# Crumpled polymer with loops recapitulates key features of chromosome organization

**DOI:** 10.1101/2022.02.01.478588

**Authors:** Kirill E. Polovnikov, Bogdan Slavov, Sergey Belan, Maxim Imakaev, Hugo B. Brandão, Leonid A. Mirny

**Affiliations:** Institut Curie, PSL Research University, Sorbonne Université, CNRS UMR3664, Paris, France; Institute of Medical Engineering and Science, Massachusetts Institute of Technology, Cambridge, MA 02139; Landau Institute for Theoretical Physics, Russian Academy of Sciences, Chernogolovka, Russia; National Research University Higher School of Economics, Faculty of Physics, Moscow, Russia; Department of Biological Engineering, Massachusetts Institute of Technology, Cambridge, MA 02142; Department of Physics, Massachusetts Institute of Technology, Cambridge, MA 02139

## Abstract

Chromosomes are exceedingly long topologically-constrained polymers compacted in a cell nucleus. We recently suggested that chromosomes are organized into loops by an active process of loop extrusion. Yet loops remain elusive to direct observations in living cells; detection and characterization of myriads of such loops is a major challenge. The lack of a tractable physical model of a polymer folded into loops limits our ability to interpret experimental data and detect loops. Here, we introduce a new physical model – a polymer folded into a sequence of loops, and solve it analytically. Our model and a simple geometrical argument show how loops affect statistics of contacts in a polymer across different scales, explaining universally observed shapes of the contact probability. Moreover, we reveal that folding into loops reduces the density of topological entanglements, a novel phenomenon we refer as “the dilution of entanglements”. Supported by simulations this finding suggests that up to ∼ 1 − 2Mb chromosomes with loops are not topologically constrained, yet become crumpled at larger scales. Our theoretical framework allows inference of loop characteristics, draws a new picture of chromosome organization, and shows how folding into loops affects topological properties of crumpled polymers.

## I. INTRODUCTION

Three-dimensional organization of chromosomes is a multi-scale complex physical system that has been challenging the field of polymer physics and stimulated the development of a broad range of polymer models. Such models mostly concern large-scale properties of chromosomes and include topologically constrained polymers [1– 4], non-equilibrium polymer states [5, 6], gels and super-coiled polymers [7], and active polymers systems [8, 9].

Recent experiments allowed characterizing chromosome folding at all scales. A Hi-C experiment produces a map of contact frequency *P* (*i, j*) between all pairs of genomic positions *i* and *j* [10]. Besides variety of local features visible in *P* (*i, j*) maps [11], the physical state of a chromosome polymer can be characterized by the scaling of the average contact probability *P* (*s*) with the genomic distance *s* = |*i* − *j*|. At large scales, in the range of *s* = 1 − 5 Mb, the scaling of *P* (*s*) ∼ *s*^−1^, markedly different from ∼ *s*^−3/2^ expected for a ideal chain (i.e. 3D random walk) or ∼ *s*^−3/2^ followed by a plateau for the equilibrium globule [12]. The *P* (*s*) ∼ *s*^−1^ scaling suggested that chromosomes are folded into the “crumpled” states [1, 10, 13] conjectured more than three decades ago [14, 15]. These polymer states are characterized by largely unknotted conformations that are stabilized by topological interactions between segments of the chain, i.e. their inability to path through each other. A chain from the melt of large unknotted non-concatenated rings is a canonical example of the topologically-stabilized polymer state, which is believed to have the fractal dimension *d*_*f*_ = 3 asymptotically (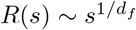 and *P* (*s*) ∼ *s*^−*γ*^, *γ* ≈ −1.1) [1, 3, 16–18] at scales *s » N*_*e*_, where *N*_*e*_ is the entangle-ment length. As we show below, existence of the minimal scale (the topological blob), at which the crumpled polymer actually develops the crumpled statistics has a crucial impact on organization of chromosomes.

At smaller scales the *P* (*s*) curve generally exhibits not a power-law behavior with a characteristic “shoulder” at *s* ≈ 100 − 200kb, which, as we show below, reflects folding of the chain into loops. We and others suggested that at smaller scales (*s* < 1 Mb), chromosomes are folded into loops formed by an active, energy-dependent, process of loop extrusion. We hypothesized that loop-extruding motors associate to a chromosome, extrude loops, and dissociate, thus maintaining the chromosome polymer in the steady state, where it is folded in an array of non-overlapping randomly positioned loops [19, 20]. Loop extrusion was observed *in vitro* [21], yet detecting and characterizing myriads of transient loops presumably present *in vivo* remains a major challenge.

Other models of interphase chromosomes folded into different types of loops, including giant rings (3 − 4Mb) [22], random overlapping loops (crosslinks) with a broad size distribution [23, 24], and rosettes of loops [25], were proposed but not systematically tested against Hi-C data. Analytical solution for even the simplest case of random crosslinks [26], however, gave *P* (*s*) ∼ *s*^−3/2^ followed by a plateau being markedly different from the experimental *P* (*s*). Dense arrays of loops were also con-sidered in models of mitotic chromosomes [27], where the interplay of factors determine the “optimal”-sized loops that maximize compaction and simultaneously minimize inter-chromosome entanglements. While accumulation of experimental data about *interphase chromosomes* continues yielding *P* (*s*) curves of similar shapes, no polymer physics model could have explained such a universal *P* (*s*), nor attempted to infer sizes and characterize organization of chromosomes.

The main challenge for an overarching model of chromosome organization is to take into account both the crumpled statistics of the chain and its folding into loops. While some attempts have been made to explain the crumpled organization itself by loops (e.g. [24, 28]), recent experiments have clearly demonstrated that they represent two independent modes of chromatin organization. Experimental depletion of loop-extruding protein complex of cohesin has removed the local shoulder on *P* (*s*) at *s <* 1Mb and revealed the power-law behavior of *P* (*s*) ∼ *s*^−1^ in the two orders of magnitude range of scales, *s* ≈ 50 − 5000kb [29, 30].

Here we develop a model of a polymer of an arbitrary fractal dimension *d*_*f*_ (including *d*_*f*_ = 3 for the crumpled state) folded into randomly positioned and non-overlapping loops, as expected to be produced out of extrusion. Our model being analytically tractable shows how such loops perturb fractal polymer organization across scales, agrees with a broad range of experimental Hi-C data, and allows to infer parameters of the loops organization. Furthermore, we reveal and describe a novel topological phenomenon, the *dilution of entanglements*, taking place in a crumpled chain folded into short-scale unentangled loops. Namely, we demonstrate that the loops would drastically increase the entanglement length *N*_*e*_ of a crumpled chromosome (∼ 10-fold or even more). As a result, a chromosome with loops turns into an unentangled though a *not* crumpled polymer at the megabase length scales. Our study provides a novel view on organization of chromosomes and, broadly, of topologically-stabilized polymer chains, where density of topological constraints can be modulated by formation of short-scale loops.

## II. A MODEL. POLYMER CHAIN FOLDED INTO LOOPS

Let us consider a classical bead-spring model of a polymer chain [12, 34, 35]. Without long-ranged interactions it corresponds to a three-dimensional random walk at sufficiently large scales (i.e. ideal chain with fractal dimension *d*_*f*_ = 2). One can generalize it to the case of arbitrary fractal dimension, *d*_*f*_ ≥ 2, via introduction of special quadratic pairwise interactions between the beads [2], in particular, allowing to describe the crumpled states with *d*_*f*_ = 3. The effective interactions result in the mean-squared spatial size of the polymer segment of *s* beads that behaves as 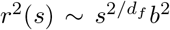 for beads of the length *b*. An important experimentally measured characteristic of such a chain is the contact probability between ends of the segment, *P*_0_(*s*), that according to the meanfield argument [1, 10, 18, 36], is inversely proportional to the volume spanned by the segment

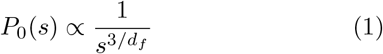

Though (Eq.1) is not true in general, it holds for the class of fractional Brownian polymers [2, 4, 37], for which (Eq.1) is the normalization of the Gaussian distribution of end-to-end distance.

Now, we consider this fractal chain folded into consecutive and non-intersecting loops with the average contour length, *λ*, and separated by gaps that have an average contour length, *g*, both exponentially distributed (Fig. 1A). Importantly, we assume that the fractal dimension *d*_*f*_ of the polymer at large scales and within a loop is not changed by the addition of the loops. Also each “loop” is modelled as an additional bond in its base, so that the ring-shaped cohesin protein does not topologically embrace two strands of chromatin but chemically binds them [38]. Then the full sequence of the gaps comprises the main chain (the backbone), controlling its equilibrium conformation properties at large scales. Clearly, such folding into loops reduces distances in polymer, turning it into a comb-like chain with loopy side bristles.

**FIG. 1.**
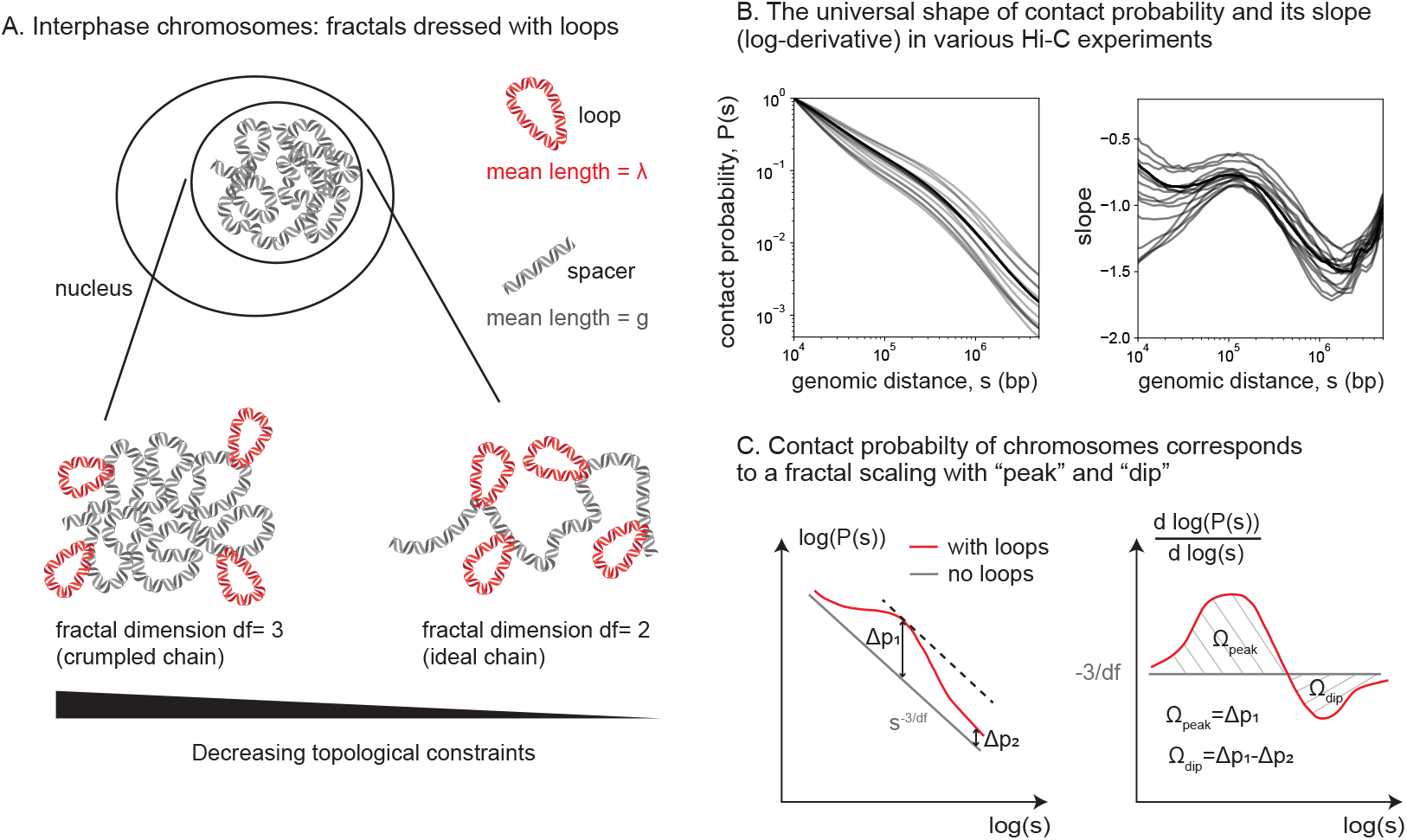
Folding of a fractal polymer into random loops recapitulates universal *P* (*s*) profiles in Hi-C experiments. **A**: a sketch of fractal polymers with different fractal dimensions *d*_*f*_ folded into loops. The mean size of the loops is *λ*, the mean size of the gaps (spacers) is *g*. **B**: a collection of 24 contact probability curves *P* (*s*) (left) and its log-derivatives (right) from experimental Hi-C and Micro-C data for different human cell types [31–33]. The shape of the profiles is universal across experimental conditions and cell types. **C**: a sketch of the typical behavior of the contact probability *P* (*s*) with and without loops (left) along with its logarithmic derivative (right) up to several megabases. A geometrical argument explains that formation of the dip is due to the difference in elevation of *P* (*s*) at small and large scales.

The effect of the loops on *P* (*s*) can be calculated using the frozen disorder approach as follows. First, one calculates the contributions of different diagrams *at equilibrium*, classifying relative positions of the points *i, j* with respect to the loop bases (Fig. 2A). For each diagram one computes the variance ⟨*r*^2^ ⟩ of the vector 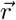 connecting the points of interest and makes use of the Gaussian relation *P* (*s*) ∼ ⟨*r*^2^⟩ ^−3/2^ for the corresponding equilibrium contact probability, see (Eq.1). For the diagrams (b) and (d) the vector 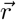 is decomposed into the sum of independent vectors 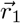 (loop), 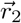 (backbone) and 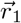 (loop), 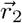 (backbone), 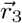 (loop), respectively. In the diagrams (b)-(d) a loop resembles a fractal bridge of the same dimension *d*_*f*_, for which the effective Hamiltonian from [2] is used (see Appendix A).

**FIG. 2.**
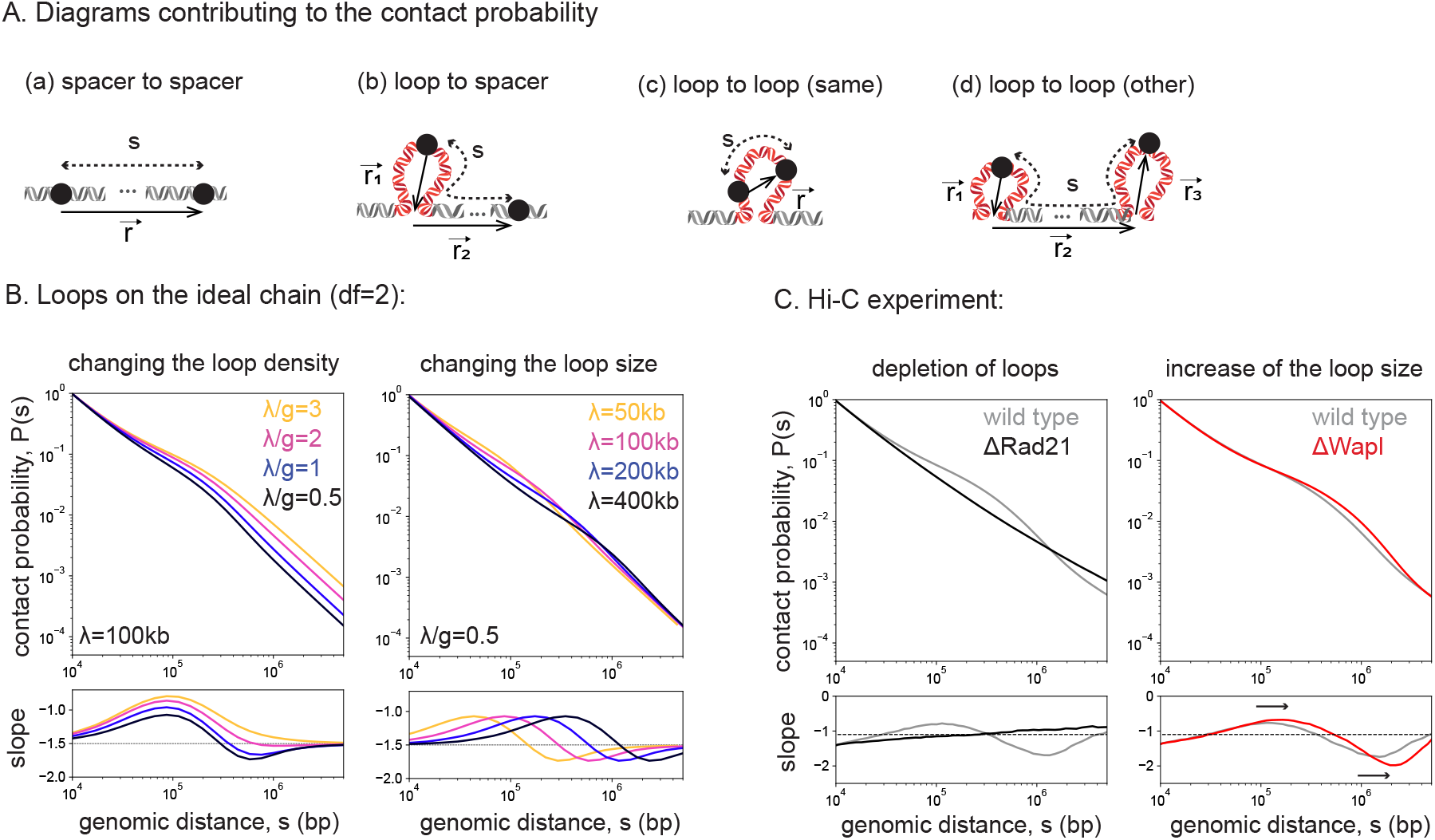
Contact probability for the fractal chain folded into an array of random loops. **A**: Illustration of the possible diagrams contributing to the contact probability. The three dots (…) denote a chromosome segment that may contain an arbitrary number of loops. **B**: Evolution of the *P* (*s*) and its log-derivative for the ideal chain (*d*_*f*_ = 2) upon the change of the parameter *d* = *λ*/*g* (left) and the mean loop length, *λ* (right), where the value of the other parameter is fixed as indicated. **C**: Behavior of the *P* (*s*) and its log-derivative in a Hi-C (specifically, Micro-C) experiment upon (left) disruption of cohesin complexes by Rad21 depletion, thus eliminating loops; (right) Wapl depletion, thus increasing of the loops size. Rad21 is a subunit of cohesin, thus, its near-complete depletion results in disruption of cohesin-mediated loops on chromosomes. Wapl protein on the contrary unloads cohesin from chromosomes, thus its depletion increases cohesin residence time and, as a result, the average loop length *λ*. The cell type is mouse embryonic stem cell, same for the two datasets [29].

Second, one averages these probabilities over all possible pairs of monomers *i, j* involving different diagrams, such that |*i* − *j*| = *s* (Fig. 2A). The exponential distribution of lengths for loops and gaps allows to make use of the well-known expression for the propagators of the two-state Markov process [39] and properly weigh contributions of different diagrams. Finally, the remaining averaging over the distribution of random loops and gaps is performed (see Appendix A). The ultimate result is factorized into the unconstrained conditional probability *P*_0_(*s*) of a loops-free chain (Eq.1) and function 𝒫 of the scaled genomic distance *s/λ* and the density parameter *d* = *λ*/*g* (for connection of this parameter to the linear loop density see [40]):

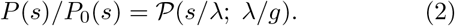

The function 𝒫 accurately accounts the contributions from four diagrams (a)-(d) depicted in (Fig. 2A); it is expressed in the form of multiple integrals involving the Bessel functions, which are to be computed numerically.

Strikingly, already for the ideal chain (*d*_*f*_ = 2) folded into loops the shape of the *P* (*s*) curve, best represented by its log-derivative, 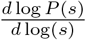, qualitatively matches the corresponding curves computed from experimental Hi-C data, see Fig. 2 B,C and Fig. 1B.

In accord with experiments (Fig. 2C), loops perturb the power-law behavior of the *P* (*s*) curve and result in the formation of a shoulder at *s* ≈ *λ*, and a corresponding “peak” and “dip” on the log-derivative plot, see Fig. 1C. Moreover, in line with our theory, experiments where loops are eliminated by depletion of cohesin (a loop-extruding motor) [29, 30] give *P* (*s*) with an almost constant slope (i.e fractal) from ≈ 50 kb to ≈ 5, 000 kb, yielding a near constant in the log-derivative without the characteristic peak and the dip (see Fig. 2C, left). Furthermore, experiments where loop sizes were increased by depleting the protein Wapl (see Fig. 2C, right) demonstrate an extension of the *P* (*s*) shoulder and displacement of the peak on the log-derivative the right. Consistently, we see that in our model (Fig. 2B) the shoulder on *P* (*s*) and the peak on the log-derivative curve, travel to the right upon the increase of the mean loop length, *λ*, at fixed *λ*/*g*.

As we see, using the simplest prior (ideal chain, *d*_*f*_ = 2) it is possible to explain qualitatively how the loops change the shape of the log-derivative. However, the ideal chain model does not capture an important aspect of chromosome organization such as the value of the *P* (*s*) slope closer to − 1, rather than 3/2 in most cell types [1, 10, 13–16]. Indeed, the ≈ − 1 scaling stretches for two orders of magnitude in genomic scale *s*, when cohesin-mediated loops are eliminated [29, 30, 41] (Fig. 2C). According to (Eq.1), it corresponds to the almost compact fractal folding with *d*_*f*_ ≈ 3. Such characteristics is a feature of the crumpled folding of a polymer, e.g. in a equilibrium melt of non-concatenated unknotted rings [1, 3, 13, 16–18] or as a long-lived non-equilibrium state of a collapsed linear chain [5, 6, 14].

The results for a non-ideal polymer with *d*_*f*_ > 2 are shown in Fig. 3A and Fig. S1. In particular, the loopy chain folded with a larger fractal dimension has a slower decaying *P* (*s*) than the ideal chain with *d*_*f*_ = 2 at all scales, the former being consistent with Hi-C data. Accordingly, the baseline (loops-free) slope of *P* (*s*), i.e. its asymptotic value 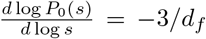, is lifting up with increase of *d*_*f*_. These properties are not surprising, since they are put in the model by construction. A non-trivial observation, however, is that the amplitudes of peak and dip on the log-derivatives notably diminish for a non-ideal chain (Fig. 3A and Fig. S1). Intuitively it can be rationalized as additional compactness (negative tangential correlations) due to loops has a weaker effect on already somewhat compact chains with *d*_*f*_ > 2.

**FIG. 3.**
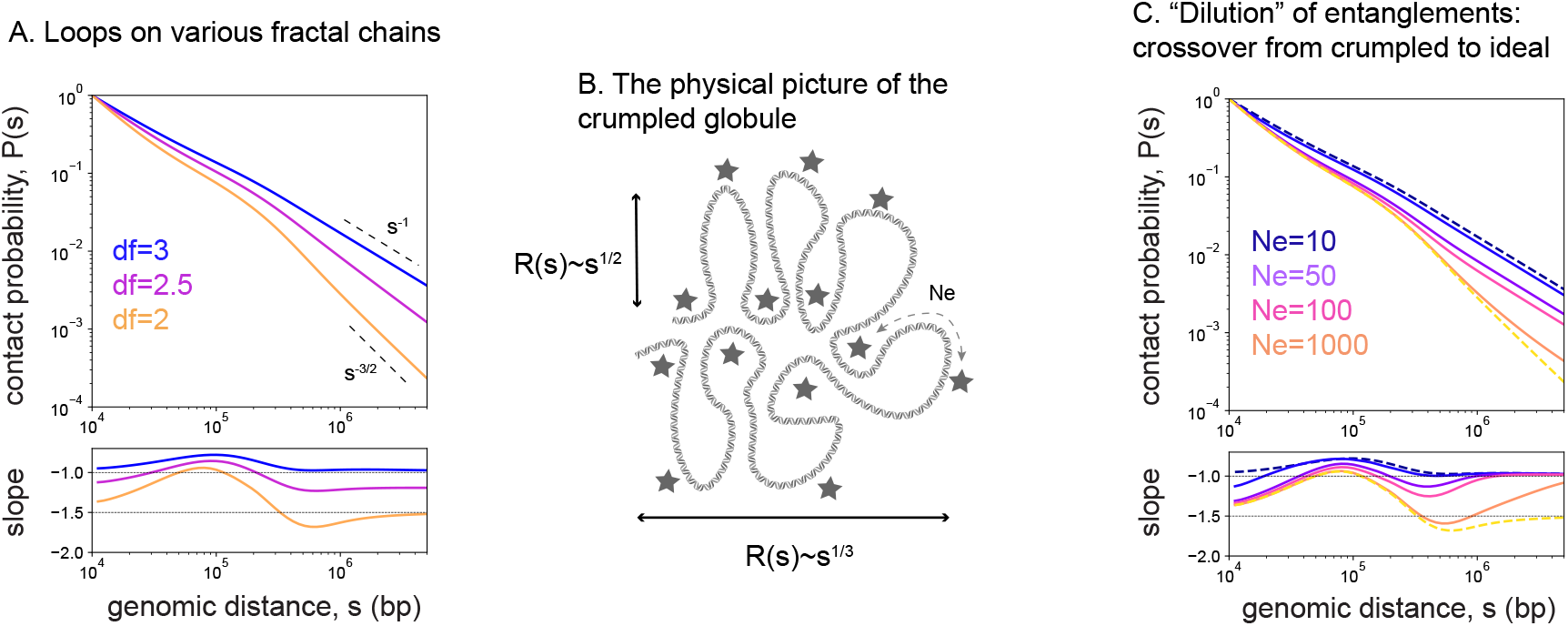
Effects in the non-ideal chain: a polymer with *d*_*f*_ > 2 folded into loops. **A**: The effect of the fractal dimension on *P* (*s*) and its log derivative (*λ*/*g* = 1, *λ* = 100kb). **B**: A physical picture of the crumpled polymer as a chain in the space with topological obstacles [42]: at short scales (between the obstacles) it is ideal (*d*_*f*_ = 2) and at large scales it is crumpled (*d*_*f*_ = 3). The entanglement length *N*_*e*_ defines a crossover length scale between the two regimes. **C**: Upon the increase of *N*_*e*_ a crumpled polymer restores ideal statistics at larger length scales (*λ*/*g* = 1, *λ* = 100kb). The dash curves respond to asymptotic loopy fractals with *N*_*e*_ = 0 (*d*_*f*_ = 3, blue) and *N*_*e*_ = ∞ (*d*_*f*_ = 2, yellow).

To additionally validate our analytical computations we compare them to numerical simulations. To calculate the contact probability *P* (*s*), (i) the loci *i, j* at the given contour distance *s* = |*i* − *j*| are sampled on the randomly generated sequences of exponentially distributed loops and gaps, (ii) the mean equilibrium distances between the loci are computed accordingly to the diagram they belong to, (iii) the distances are translated into the contact probabilities using the mean-field relation (Eq.1) and, finally, (iv) the sample averaging is performed. This approach essentially allows to weight the contributions of different diagrams numerically via statistical sampling, while keeping the end-to-end distribution Gaussian. As we demonstrate in the Fig. S3, the numerical approach perfectly agrees with the results of our theory, irrespective of the value of *λ*/*g*.

A simple argument helps to understand how loops perturb the contact probability *P* (*s*), generating the curves we observe in the theory and experiments. Indeed, the loops impose two effects on *P* (*s*) at different scales. At small scales, the dominant contribution to the elevation of *P* (*s*) comes mainly from contacts between two loci within the same loop (e.g. see Fig. S2, for *λ*/*g* ≥ 1/5). Within a loop, the contact probability is increased due to the smaller physical size of a loop compared to an open chain of the same contour length, and due to a shorter effective contour length between two points near the loop base. Thus,

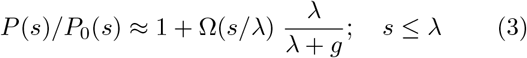

where Ω(*s*/*λ*) is some function describing the shape of the shoulder, which is weighted by the fraction of the chain within the loops.

At large scales, when *s* » *λ*, the contact probability becomes elevated due to a shorter effective contour length between the two points, since the shortest path between monomers skips all intervening loops:

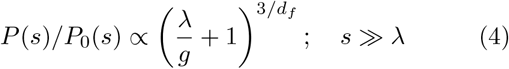

The interplay between the shoulder at small scales (Eq.3) and the elevation of *P* (*s*) at large scales (Eq.4) determines the shape of *P* (*s*) as well as the “peak” and the “dip” of the log-derivative. The peak is always present as a result of breaking of the scale-invariance due to loops formation. Fig. 1C displays a simple geometric condition for the dip: the shoulder, (Eq.3), at *s ≈ λ* must be higher than the elevation at *s » λ*, (Eq.4). As the spacer, *g*, shrinks (loop density increases), the elevation at large scales (Eq.4) indefinitely grows, while the height of the shoulder (Eq.3) eventually saturates. Therefore, upon a gradual compaction of the chain into an array of loops, the dip becomes more shallow and dissolves. As the solution for the ideal chain (*d*_*f*_ = 2) suggests (Fig. 2B, left, Fig. S1), there is a critical value of parameter *d* = (*λ*/*g*)^∗^ ≈ 2, at which the dip completely disappears. For the compact chain (*d*_*f*_ = 3) the corresponding critical values is lower, *d* = (*λ*/*g*)^∗^ ≈ 1. Indeed, since the absolute perturbation at the shoulder Ω(*s/λ*) is smaller for chains with *d*_*f*_ > 2, the dip disappears at a lower loop density than for the ideal chain. Physically, this marks a crossover from sparse to relatively dense array of loops where the effective shortening of the polymer at large scales reaches the level of compaction at scales of a single loop.

### Loops on a crumpled chain

The crumpled statistics cannot be realized at all scales in a physical system [3, 14, 16, 17]. For a physical crumpled chain there is a crossover length scale *N*_*e*_, called the entanglement length (having the meaning of the topological blob), only above which the compact folding with *d*_*f*_ = 3 can take place. Below this scale the topological constraints are not sufficient and the segment remains ideal with *d*_*f*_ = 2 down to the persistent length (for a loosely entangled chain) or the concentration blob size [12, 34]. The end-to-end squared size of a segment *s* of a crumpled chain with the entanglement length *N*_*e*_ and the persistent length *l*_*p*_ can be written as

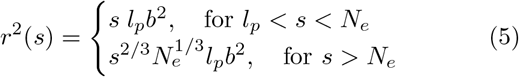

To quantitatively explore the effect of this crossover in real crumpled chains on *P* (*s*), we generalize our theory to a *non-fractal* chain governed by (Eq.5). For smoothness of resulting *P* (*s*) we allow the crossover point in (Eq.5) to fluctuate around the mean *N*_*e*_ within the exponential distribution (see Appendix A).

We find that considering the scale *N*_*e*_, where topological effects start to play important role, changes the behavior of the *P* (*s*) in the model. With increasing *N*_*e*_, the crumpled polymer with loops starts to show much more pronounced “peak-and-dip” on the log-derivative curve (Fig. 3C). Since for *s* < *N*_*e*_ the system behaves like an ideal chain with loops, for *N*_*e*_ *» λ* the effect of topological crumpling becomes irrelevant and the log-derivative curve resembles that of loops on an ideal chain (Fig 3A, *d*_*f*_ = 2). Fig. 3C demonstrates this crossover from the crumpled (blue) to ideal chain regimes (yellow), upon the increase of *N*_*e*_, with a pronounced “peak-and-dip” emerging when *N*_*e*_ ≈ *λ*. For example, for *λ*/*g* = 1, and *N*_*e*_ = 50 ∼ *λ* = 100, not atypical for dense polymer systems, one has a pronounced dip on the log-derivative, which is otherwise absent in the model without *N*_*e*_. Further increase in *N*_*e*_ results in even deeper dip, while the amplitude of the peak located at ≈ *λ* saturates.

The pronounced dip requires *N*_*e*_ ≳ *λ*, which in turn indicates that extruded loops cannot be crumpled. For a ring to be crumpled it needs to be several dozens of *N*_*e*_ in length (10 − 30*N*_*e*_) [18, 43]. At the same time, the entanglement length for chromatin, estimated in the literature (see Table 2 in [1]; *N*_*e*_ ≈ 20 − 100kb) is comparable to the size of cohesin-mediated loops *λ* ≈ 100 − 150kb. Therefore, this picture suggests that the extruded loops in chromatin are practically not crumpled (unentangled).

Remarkably, a crumpled chain with unentangled loops captures both the pronounced peak and dip and levelling of the curve around ≈ − 1 both seen across Hi-C experiments, and not captured by models that we considered above (the ideal chain, and the fractal chains with *d*_*f*_ > 2).

### Dilution of entanglements in a crumpled chain induced by folding into loops

Can folding into unentangled loops, in turn, affect entanglements of the main chain? In what follows we suggest that since the loops are unentangled they do not impose any considerable topological constraints for the main chain. As more material goes into loops, the main chain shortens, which leads to the decrease of the total amount of entanglements (dilution) and increase of *N*_*e*_ of the main chain. This effect, that we refer to as the “dilution of entanglements”, can be physically understood as screening of entanglements by the loops inside the growing topological blob. As we demonstrate below, *N*_*e*_ of the main chain indeed grows with the loop density, that in turn affects the contact probability *P* (*s*) of the loopy chain across scales. This way effects of large *N*_*e*_ considered in the previous section becomes even stronger.

To estimate the effect of loops on *N*_*e*_ of the main chain we start with a known empirical expression for dense linear chains [1, 44, 45]:

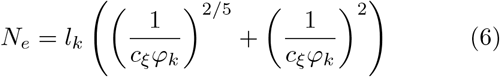

where *l*_*k*_ is the Kuhn length (in monomers), 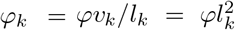 is the dimensionless volume density of Kuhn segments of volume *v*_*k*_ and monomer volume density *φ*; *c*_*ξ*_ ≈ 0.06 is a phenomenological constant that was used to describe various simulation data for entangled polymers [44]. For loosely entangled linear chains, *φ*_*k*_ < 1, the quadratic factor in (Eq.6) dominates. Furthermore, for crumpled territoriral chains described by (Eq.5) one can straightforwardly show that the Kuhn volume density of a single chain is similarly related to its entanglement length,

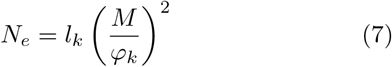

for all *φ*_*k*_ and where *M* ∼ *O*(1) is the number of other chains populating the volume of a marked crumpled chain. The dependence of *N*_*e*_ on *φ*_*k*_ for the crumpled polymer can be understood in the picture of a chain in the array of topological obstacles (constraints) [46, 47], which is collectively generated by spatial contacts between the crumples (see Fig. 3B, Fig. 4A). Clearly, not all spatial contacts contribute to topological obstacles. However, similarly to linear chains [45], non-specific dilution of the contacts upon decrease of *φ*_*k*_ leads to a fewer topological obstacles in the system and, accordingly, increase the entanglement length, *N*_*e*_ (Eq.7).

**FIG. 4.**
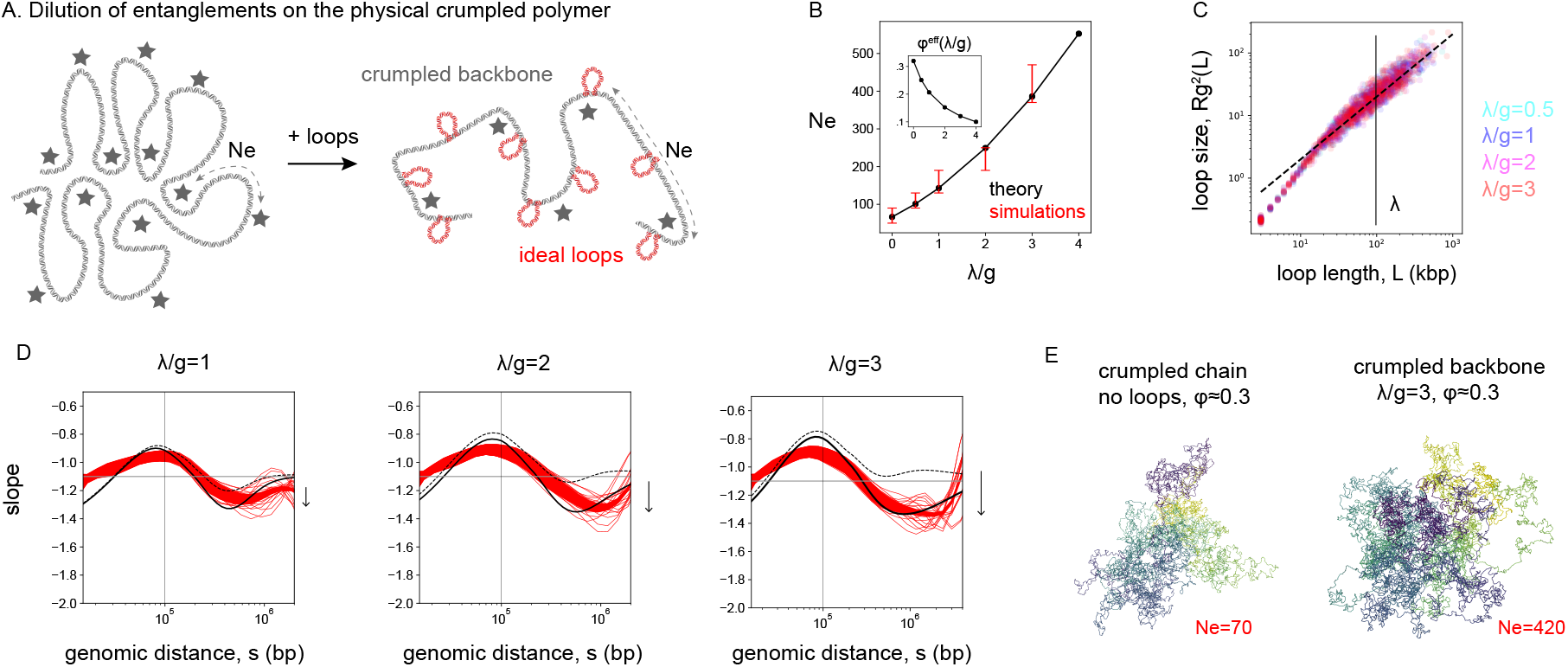
Dilution of entanglements upon folding of a crumpled chain into loops. **A**: A sketch for the effect of dilution of entanglements upon addition of the loops. Short-scale loops are unentangled and do not contribute to the crumpling of the backbone. Shortening of the backbone of the chain with loops (right) as compared to the loops-less situation (left) drives reduction of entanglements and increase of *N*_*e*_. **B**: Increase of the entanglement length as a function of parameter *d* = *λ*/*g* as measured in simulations (red) and computed in the theory according to (Eq.6)-(Eq.8) (black). For the theoretical curve the conventional value of the parameter *c*_*ξ*_ = 0.06 in (Eq.6) is used, which was previously found in [44] to match various entangled systems; the value of the microscopic constant *c*_*α*_ in (Eq.8) is found to be 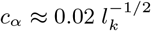, which yields *c*_*α*_ ≈ 0.01 for *l*_*k*_ ≈ 4 kb. The corresponding decrease of the effective monomer volume density 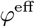 (which is 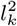 times smaller than 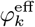 in (Eq.8)) is shown in the inset. **C**: Squared gyration size of the loops in simulations as functions of their contour length for various loop densities (*λ*/*g* = 0.5, 1, 2, 3). The average loop size is *λ* = 100kb. **D**: Slopes of the contact probability *P* (*s*) for various densities of the loops *λ*/*g* = 1, 2 and *λ*/*g* = 3 from simulations (red); theory with fixed *N*_*e*_ = *N*_*e*_(*λ*/*g* = 0) = 70 (dashed black); theory with *N*_*e*_(*λ*/*g*) that is growing with the loop density, according to Fig. 4B (solid black). The value *d*_*f*_ = 2.7 (not *d*_*f*_ = 3) is chosen for the theory as it corresponds to the loops-free slope 3*/d*_*f*_ ≈ 1.1 in simulations (not − 1). The red strip of the simulations is bound by two values of cutoff (capture radius of contact): *r*_*c*_ = 3kb and *r*_*c*_ = 5kb. Different thin red lines correspond to different replicates. The average loop size is indicated by the vertical line, *λ* = 100kb. The arrow indicates deepening of the dip in the theory upon taking into account the dilution of entanglements. **E**: The snapshots from simulations are shown for the backbone of a chain with loops (*λ*/*g* = 3, right) and for the main chain with cut and removed loops and further confined to the original monomer volume density *φ* ≈ 0.3 (left).

We suggest that due to the commutative nature of topological constraints, unentangled side loops do not contribute to creating of the topological obstacles and hence do not crumple the main chain. Thus, in our sys-tem the *effective* volume density 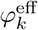, relevant for the entanglements, is controlled by the backbone, which is shortening upon folding into loops. Still the loops occupy a certain excluded volume in the space, creating the crowding effect for the main chain. The two phenomena, backbone shortening and loops crowding, are encoded in the theory for 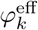 of a crumpled polymer folded into loops of size *λ* and density parameter *d* = *λ*/*g* (see the full derivation in Appendix B):

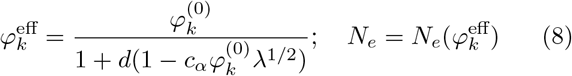

where the constant *c*_*α*_ depends on the microscopic struc-ture of the fiber, 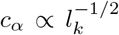; we treat *c*_*α*_ as a free parameter in our model. The structure of (Eq.8) is quite transparent. Without loops (*d* = 0), the backbone resembles the original crumpled chain with effective volume density 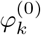. If the loops are sufficiently short so that their volume can be neglected, the decreased effective volume density is fully described by the backbone shortening, which yields the factor *d* + 1 = (*λ* + *g*)*g*^−1^ in the denominator of (Eq.8). In general, the loops partially compensate this effect by crowding the own volume of the chain, which leads to an increase of 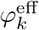. Importantly, as we show in the Appendix B, the compensation by the loops is always not complete 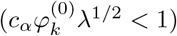 for unentangled loops.

The effect of dilution of entanglements (Eq.6)-(Eq.8) on *P* (*s*) can be well-captured by extending our analytical framework (the full theory). For that we consider ideal loops, while the entanglement length *N*_*e*_ of the crumpled backbone is now a certain function of loop size and loop density, 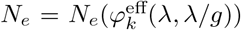. Qualitatively, the effect on *P* (*s*) can be understood as follows. As previously, the shape of *P* (*s*) at scales *s* ≈ *λ* (Eq.3) is largely determined by intra-loop contacts and is not sensitive to organization of the backbone chain. However, at large scales *s* » *λ* the contact probability is now additionally decreased by the factor of 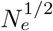, reflecting spatial segregation of the loops due to the dilution of entanglements of the main chain

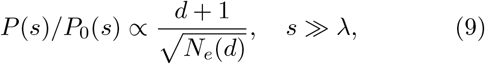

As we see from (Eq.9), the increase in contact frequency due to chain shortening is now counteracted by spatial separation of the loops due to the increased *N*_*e*_ (Fig. 4A). Since in the regime of interest the entanglement length quadratically depends on the density parameter *d*, as seen from (Eq.7) and (Eq.8), at *s* » *λ* one has

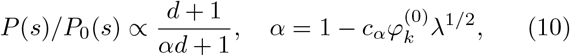

Interestingly, for sufficiently short unentangled loops, *α* ≈ 1, the backbone decompaction *almost fully compensates* its shortening (the crowding of loops, i.e. *α* < 1, still slightly elevates *P* (*s*) at small *d*), see Fig. 5A and Fig. S8. This result is physically related with the territorial organization of crumpled chains. Indeed, the size of the crumpled chain is controlled by the combination 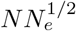, see (Eq.5); shortening of the backbone due to loops by a factor *d* + 1 leads to increase of 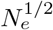 by the same factor. As a result, crumpled chains with short-scale loops tend to decompact their shortened backbones to the volume of their own territories, nearly maintaining the spatial organization at large scales. While at large scales *P* (*s*) is almost unchanged (Δ*p*_2_ = 0 in Fig 1C), at short scales it is elevated due to folding into loops (Δ*p*_1_ > 0), see (Eq.3), which ensures formation of the dip independently of the loop density (Fig. 1C). Thus, territoriality of crumpled chains underlies the observed universality of experimental *P* (*s*) curves (Fig. 1B).

**FIG. 5.**
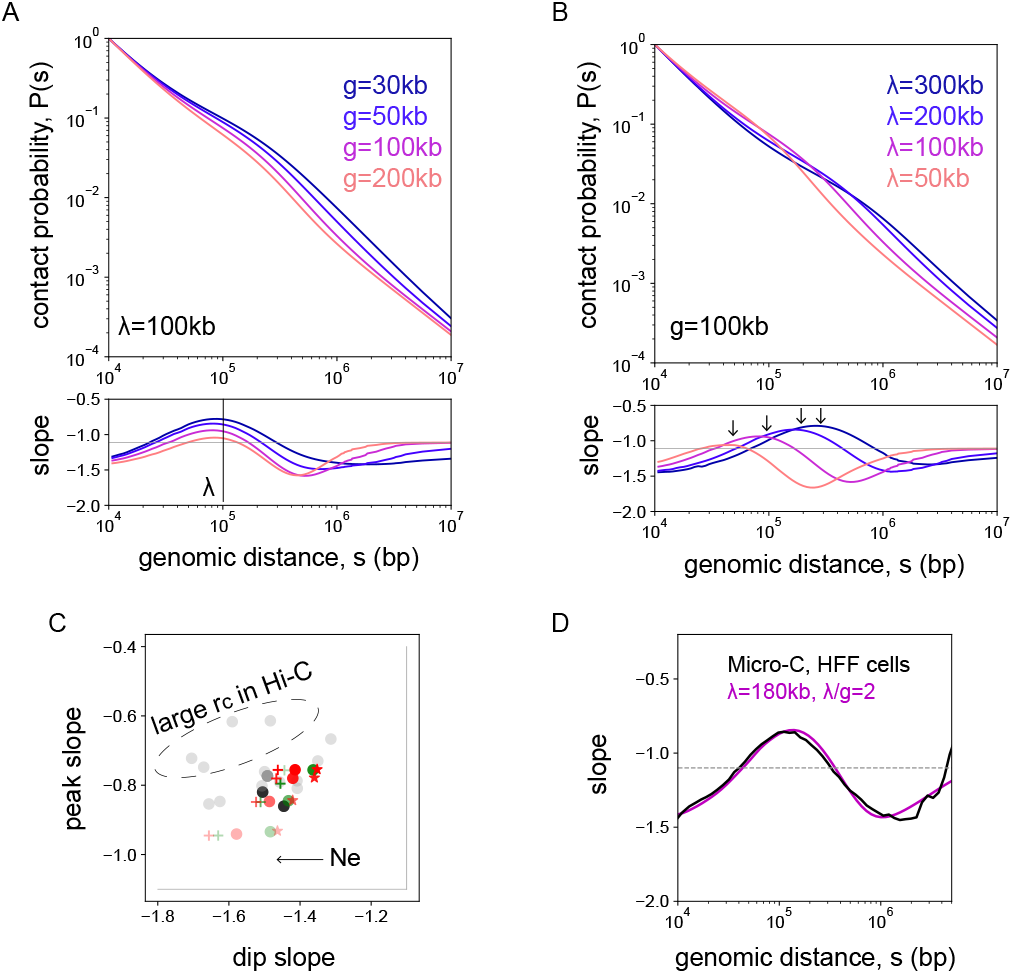
Behavior of the contact probability curves in the full theory with ideal loops and calibrated *N*_*e*_(*λ*/*g*). **A**: Change of spacer size, *g*, while loop size is fixed, *λ* = 100kb. **B**: Change of loop size, *λ*, while spacer size is fixed, *g* = 100kb. Loop density increases from red to blue color. **C**: Diagram of slopes at the peak and at the dip for various theoretical and experimental curves. Experiment: Hi-C (gray), Micro-C (black); Theory: *λ* = 100 (red), *λ* = 200 (green); *φ* = 0.1 (crosses), *φ* = 0.2 (circles), *φ* = 0.3 (stars); the dots get less transparent with increase of the loop density. **D**: Fit of computed experimental Micro-C log-derivative (data from [32]) by the theory with *λ* = 180kb, *λ*/*g* = 2 and volume density *φ* = 0.2. The resulting value of the entanglement length is *N*_*e*_ ≈ 700, i.e. at the scale of ≈ 8 loops the main chain is unentangled.

Taken together, the phenomenon of dilution of entanglements in real crumpled chains results in significant increase of *N*_*e*_ of the main chain upon folding of a polymer into loops. This effect in turn restores the dip on the log-derivative (Fig. 4D), compared to the model with fixed *N*_*e*_, and generates the universal shapes of the contact probability observed experimentally.

#### Simulations of crumpled chains with loops

Next we examine the dilution of entanglements by direct polymer simulations. To test predictions of theoretical argument above we perform equilibrium polymer simulations of a crumpled loopy polymer rings (*N* = 90000 beads, ≈ 90Mb chromosomal region) with excluded volume and topological constraints (see Appendix C).

We observe that despite of topological constraints, and consistent with the theory, the loops on the chains are indeed ideal, as their squared gyration size 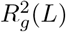 grows linearly with their contour length, *L*, Fig. 4C.

Furthermore, we observe the increase of entanglement length *N*_*e*_ of the backbone with *d* = *λ*/*g*. In order to accurately check for this effect, we infer *N*_*e*_ and *l*_*p*_ by fitting *R*(*s*) of the backbones from simulations by a theoretical curve. This curve was constructed by extending the worm-like chain model [48] to take into account the crossover from ideal to crumpled statistics at *N*_*e*_ (see Appendix C for details). Importantly, while the persistence length of the chain remains unchanged (*l*_*p*_ = 1.75) as a result of addition of loops, we see that the entanglement length grows 6-fold from *N*_*e*_ ≈ 70 at *d* = 0 to *N*_*e*_ ≈ 420 at *d* = 3 (Fig. S4). As we find, the values of entanglement length of the backbone directly measured in simulations are in excellent agreement with our theoretical predictions by (Eq.6) and (Eq.8), see Fig. 4B.

Next we examined the physical origin of the backbone decompaction, i.e. the growth of *N*_*e*_. The theory suggests that this effect is due to dilution of entanglements, and not due to the extension of the backbone by repulsion between the loops. To test this, we run additional simulations where we cut off loops from the backbone, turning it into the system of disconnected loops and the backbone. Then we equilibrated the resulting system under the same total volume density. First, we observe that the loops remain Gaussian, despite they are now deattached from the backbone (Fig. S5 A), and their sizes are unchanged. Second, for the same loop density the system with cut loops and the original one with attached loops produce indistinguishable, within the error bar, *R*(*s*) of the main chain (Fig. S5 B, C). We conclude that loop attachment and interactions between them do not deform the main chain.

While occupying some volume, loops do not crumple the main chain either. In fact, when we confined the single backbone chain to the same volume density, but without any loops, it became more crumpled, yielding a smaller *N*_*e*_. Indeed, the effective substitution of the unentangled loops by the crumpled backbone in a unit volume results in stronger crumpling and the decrease of the entanglement length (Fig. S6, Fig. 4E). The observed value of *N*_*e*_ corresponds to the loops-free system, *N*_*e*_ ≈ 70. These results demonstrate that the loops indeed induce topological screening and decompaction of the main chain, however, their attachment to the backbone plays no role in this effect.

Simulations confirm both the increase in *N*_*e*_ and the mechanism of “dilution of entanglements” put forward by our theory. The main factor driving this effect is shortening of the main chain, accompanying by screening of entanglements by the unentangled loops (Eq.8).

In good agreement with the theoretical argument (Eq.9) in polymer simulations we further find that the shapes of *P* (*s*) log-derivatives demonstrate a pronounced amplitude irrespective of the loop density (Fig. 4D). As was explained above, this is a signature of increasing *N*_*e*_ of the backbone. In Fig. 4D we compare the log-derivatives of *P* (*s*) for the simulations of crumpled chains folded into loops with the theoretical curves computed for the original *N*_*e*_ = 70 (no increase of *N*_*e*_ induced by the loops), corresponding to the loops-less case (dashed black curves). It is clear that the model with fixed entanglement length cannot explain pronounced dip at all loop densities. On contrary, as the theory suggests, the increase of *λ*/*g* leads to gradual vanishing of the dip (Fig. 2B, Fig. S1), as long as *N*_*e*_ is fixed.

When we calibrate *N*_*e*_ according to the value of *λ*/*g* (Fig. 4B) and compute the corresponding theoretical *P* (*s*) log-derivatives, we see the restoration of the dip and much better quantitative agreement with simulations (solid black curve in Fig. 4D). Still the agreement is not perfect, since the theory is Gaussian, while crumpled polymers are clearly not [36], which is responsible for subtle deviations between the theory and simulations. Despite of this, we conclude that the theory with calibrated entanglement length *N*_*e*_ = *N*_*e*_(*λ*/*g*) quantitatively accounts for the behavior of crumpled chains with loops in simulations and restores the dip on the log-derivatives. We thus emphasize that the observed effect of dilution of entanglements in crumpled chains leaves distinct signatures in the *P* (*s*) curves – pronounced peak and dip, and leveling at − 1 slope – which is evident in the Hi-C data for a broad range of cells and conditions.

## IV. DISCUSSION

A polymer folded into loops is a new and exciting physical system. Surprisingly, loops affect not only local fold-ing of the polymer, but also long-range organization of the chain, and its topological characteristics. As we show, the interplay between the loops-induced compaction at small scales and arrangement of an array of loops at large scales results in a characteristic shape of the *P* (*s*) seen across cell types and organisms. We find that the entanglement length *N*_*e*_ in crumpled chains and the comb-like organization of a loopy chain lead to an interesting effect where the loops and the main chain at the scale of a few loops are not crumpled. Moreover, folding into loops further reduces the density of topological entanglements, a phenomenon we refer as the *dilution of entanglements*. All these effects together lead to the shape of the *P* (*s*) log-derivative curve, similar to experimental one: with a pronounced peak and dip. Ultimately, this agreement with experiments suggests that chromosomes are folded into loops that are not crumpled (unentangled), and neither is the main chain at the scale of a few loops (∼ 1 − 2Mb). Yet, at larger scales (> 10 loops) the main chain starts forming a crumpled object. This result is in striking diversion from the current understanding of chromosome organization.

Importantly, as we further show in simulations (Fig. S7 A,B), compartmentalization of chromosomes does not affect the characteristic contact probability curve of a polymer folded into loops. Indeed, the typical scale of compartmental domains (in humans) is several megabases, which is an order of magnitude larger than the loop size, *λ*. When block-copolymer interactions derived from experimental Hi-C maps (see Appendix C) are added into simulations to the fixed loops, the resulting *P* (*s*) curves do not change significantly (Fig. S7B); the constant ≈ − 1.1 slope of the loops-free crumpled chain is also unperturbed by addition of compartments (Fig. S7A). Our result is also consistent with experiments where depletion of cohesin caused expected changes in the *P* (*s*) curves (the loss of the peak and dip on the log-derivative), yet retention and strengthening of compartments (Fig. 2C) [30, 41].

The dilution of entanglements is a novel exciting phenomenon in crumpled chains folded into loops. We suggest that formation of loops shortens the main chain and stores the material inside the unentangled loops; if loops are short (*λ* ≈ *N*_*e*_), they are neither crumpled nor can crumple the main chain, serving as a “reservoir” of unentangled material. This evidently leads to reduction and dilution of topological obstacles along the main chain, both globally (the total amount of the obstacles) and locally (linear density of the obstacles). The latter results in the increase of entanglement length *N*_*e*_ of the main chain and, therefore, decompaction of chromosomes, which counteracts the effective contour length shortening. We suggest a theory explaining how this dilution effect depends on parameters of the loops, which is supported by the simulations. A similar effective stiff-ening of chromosomes has been recently suggested experimentally [49].

The developed theory allows to rationalize the shapes of experimental curves in different cell types and upon the change of conditions. Our model shows how sizes of loops, not directly visible in Hi-C data, can be inferred from the shape of experimental *P* (*s*) curves. First, the peak on the log-derivative corresponds to the size of the loops with a factor of order of unity (Fig. 5 A,B, Fig. S8), marginally dependent on the loop density. This observation allows to estimate the typical sizes of chromosome loops from *in vivo* data, yielding a range of *λ* = 100 − 200 kb for various human cells (Fig. 1B). The diagram of *P* (*s*) slopes at the peak and the dip further allows to infer the typical values of density parameter *d* and loop sizes *λ* from comparison of experimental data points with our theory (Fig. 5C). However, such an analysis of Hi-C curves should be done with caution: it is known that the coarse capture radius in Hi-C can flatten the *P* (*s*) at short scales (raising its log-derivative) [32, 33] (see also Fig. 1B). Accordingly, we see in Fig. 5C that a few Hi-C data points have larger amplitudes of the peak slope than the theory can explain. At the same time we find a rather good agreement of our theory with Micro-C experiment, where the capture radius is lower, approaching the size of a single nucleosome. The fit presented in Fig. 5D suggests that the loop size is *λ* ≈ 180kb, which also corresponds to the value derived from the peak position, and *λ*/*g* = 2.

These estimates draw a picture of chromosome organization very different from a commonly accepted view (Fig. 6). For the estimated *λ* and *λ*/*g*, the entanglement length (of the main chain) becomes significantly large, *N*_*e*_ ≈ 700kb, further suggesting that at scale of *N*_*e*_*/g* ≈ 8 loops or *s* ≈ 2Mb of genomic distance, a chromosome is virtually unentangled. This result is a clear manifestation of the dilution of entanglements due to folding into loops. At *s* < 2Mb a chromosome backbone follows the statistics of an ideal chain (not crumpled), while at larger scales it becomes crumpled. When the loops are removed, however, *N*_*e*_ drops to *N*_*e*_ ∼ 50 − 70kb [1] and the chain becomes crumpled in a large range of scales, experimentally 50kb< *s* < 5000kb [29, 30] (Fig. 2C).

**FIG. 6.**
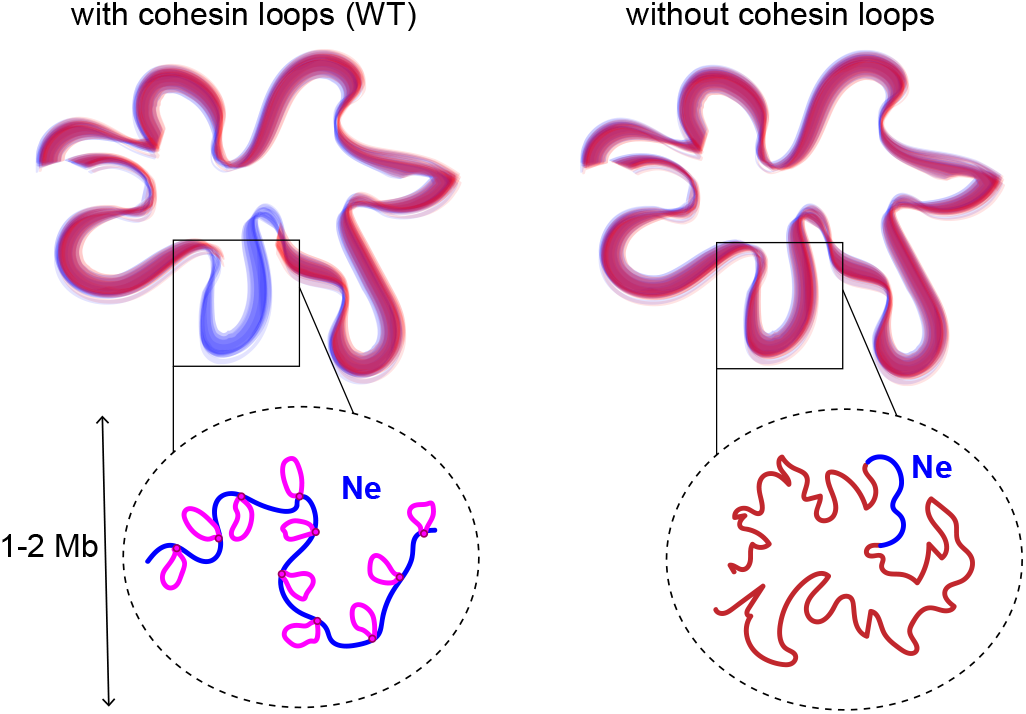
Chromosomes organization with (left) and without (right) cohesin loops. At large scales *s* > *N*_*e*_ the chains are crumpled, and at smaller scales *s* < *N*_*e*_ they remain ideal. Dilution of entanglements due to loops leads to increase *N*_*e*_ ≈ 1 − 2Mb, compared to the loops-free case when *N*_*e*_ drops to *N*_*e*_ ≈ 50 − 70kb.

Our theory has several implications and makes testable predictions. Unentangled organization of chromosomes at submegabase scales may be consistent with chromosome dynamics that appear to have Rouse scaling of MSD(*t*) ∼ *t*^1/2^ [50]. Our theory also suggests that tracing of the backbone (between the loops) in microscopy, if possible, would produce ideal statistics of the intrachain distances *R*(*s*) ∼ *s*^1/2^ up to several Mb with loops (Fig S4), while ∼ *s*^1/3^ without loops (as seen in [51]). More sparse entanglements in chromosomes with loops can also potentially render different mechanical properties of cell nuclei as compared to the case without loops. In Polymer Physics, we expect to observe a similar phenomenon of dilution of entanglements in equilibrium melt of linear chains folded into loops, where it can affect dynamics and response to mechanical stress.

While being analytically tractable and powerful in explaining experimental data, our theory has several limitations, that we outline below.

First, we consider here independent and exponentially distributed loops and gaps. This allows us to map the sequence of loops and gaps onto the two-state Markov process and obtain a precise expression for the weights of the diagrams. In the dynamic model of loop extrusion there exist nested loops, correlations between the consecutive loop and gap sizes and stalling interactions (collisions) between the neighboring motors, as well as with other proteins on chromosomes. It was shown before [20] that abundant collisions can result in a distribution of the loop sizes which can be approximated by the normal. However, the analytical form of the distribution of gaps for the simplest case of non-interacting (sparse) motors, as well as the microscopic rules of real extrusion in cell are not known at the present time.

Second, we consider only equilibrium loops, whereas loops actively extruded by motor proteins turn a polymer into an active/non-equilibrium system. The fixed loops approach relies on the assumption that the time to actively extrude the loop is larger than the time of its passive relaxation. Thus, there exists a critical length scale, *s*^∗^, such that for loops *λ* < *s*^∗^ the relaxation is faster than extrusion, and our assumption largely holds. Quan-titative estimates (see Appendix D) indicate that for the upper limit of the extrusion speed of *r* = 1 kb/s [52], the Kuhn segment *l*_*k*_ = 4kb and Rouse diffusion coefficient *D*_*R*_ = 10^−2^*μm*^2^*s*^−1/2^ [50] the critical equilibrium scale can be as large as *s*^∗^ = 100 *÷* 1000kb 2: *λ*. Another experimental estimate from [50] is that a chain of ∼ 500kb equilibrates in ∼ 40min; extrapolating to 100kb loops it gives ∼ 2min to equilibrate. Loops of *λ* ≈ 100kb can be extruded in as little as ∼ 2min if the speed is 1kb/sec or in ∼ 10min if experimentally measured cohesin residence time is used. Thus, at the lower limit of this range, loops may not be fully equilibrated. We note that equilibration of the loops and restoration/extinction dynamics of topological obstacles in the steady-state would create a non-trivial interplay, whose consequences on the contact probability are yet to be understood.

Third, for the sake of analytical tractability we employ here a Gaussian model, mapping a fractal chain onto the fractional Brownian motion [2]. We explicitly rely on the Gaussianity of the chain by making use of the meanfield connection between the equilibrium spatial distance and the contact probability (Eq.1). Moreover, the spatial distances in loops were computed by treating them as fBm bridges, which allowed us to exploit the effective Hamiltonian of fractal polymer states suggested recently [2, 37].

Fourth, we do not consider very dense arrays of loops, bottle brushes, that fold chromosomes into elongated compacted bodies (during mitosis [53], meiosis [54], and WaplKO). At *λ/g >* 4, the size of a Gaussian loop reaches the size of the Gaussian gap, marking a transition to the bottle-brush regime, when rigidity of the chain can become dominant. As shown in [55] for comblike chains with linear side groups, in this regime (specifically, a loosely grafted bottle-brush) the gaps become fully stretched due to osmotic pressure generated by the side chains. In case of crumpled backbone a non-trivial interplay between the stretching and decompaction (increase of *N*_*e*_) can happen in the dense regime. Consistently the bottle-brush stiffening might be responsible for a slight deepening of the dip observed in Wapl knock-out experiments (Fig. 2C), where the increased abundance and processivity of cohesin (smaller *g* and larger *λ*) might turn a chromosome in a dense mitotic-like bottle brush.

In summary, we have developed an analytical theory of a crumpled polymer folded into loops and have revealed a novel topological phenomenon in such a system. We demonstrate that information about loops is contained in the shape of experimentally measurable contact probability curve, *P* (*s*), and primarily in its log-derivative. Our theory quantitatively reproduces the experimental curves and allows to infer parameters of the loops *in vivo*. Furthermore, we propose that the density of entanglements in chromosomes is reduced due to folding into loops, resulting in chromosomes that are *not* crumpled at scales up to ∼ 1 − 2 Mb. Together, our findings demonstrate that folding of a polymer into loops not only changes its conformational characteristics across scales, but also reduces the topological constraints in the polymer system.

## ACKNOWLEDGEMENTS

We thank Mehran Kardar, Alexander Grosberg, Ralf Everaers and Mikhail Tamm for valuable discussions on the subject of the paper. LM acknowledges support from the NIH Common Fund 4D Nucleome Program, HG011536, Center for 3D Structure and Physics of the Genome, and NHGMS NIH R01GM114190. The work of KP and BS is supported by the Russian Science Foundation (Grant No. 21-73-00176). KP also acknowledges the support of the Programme d’investissements d’avenir (LabEx DEEP). The work of SB on the ideal chain model has been supported by the Russian Science Foundation (Grant No. 20-72-00170). SB also acknowledges support from James S. McDonnell Foundation in 2017-2020.

## APPENDIX A

### DERIVATION OF THE CONTACT PROBABILITY FOR A FRACTAL POLYMER CHAIN WITH LOOPS

#### 1. Basic properties of equilibrium ideal Gaussian chains

In the absence of SMC-mediated loops, the probability distribution of the separation vector 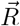 between two sites of an equilibrium ideal linear Gaussian chain separated by contour distance *s* is given by (see Ref. [12, 34, 35])

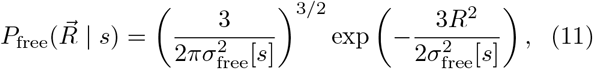

where

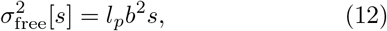

and *b*^2^ is the mean square length of a chain segment, *l*_*p*_ is the persistence length. Gaussian property is supposed to be if *s » l*_*p*_.

For an equilibrium ideal Gaussian bridge having size *L*, the probability distribution of the separation vector 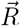 between two sites separated by contour distance *s* (< *L*) is given by

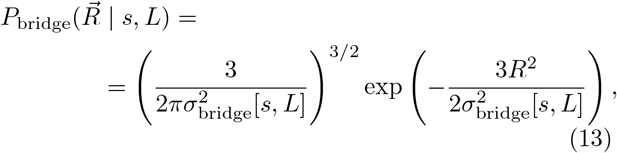

where (see [12])

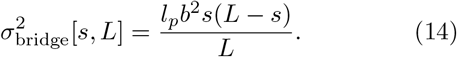

#### 2. Subdiffusive fBm trajectories and the effective Hamiltonian

To take into account non-Markovian properties of the trajectory at values of fractal dimension *d*_*f*_ > 2, we consider the class of fractional Brownian motion (fBm) with the Hurst parameter *H* = 1*/d*_*f*_ as a model to the fractal polymer fiber. To begin with, define the trajectory of a discretized random walk, or, similarly, a conformation of a polymer chain in the three-dimensional space by a set of coordinates {***X***} = {***x***_0_, ***x***_1_, …, ***x***_*N*_}. There exist a unique stationary measure *P* (***X***) over the realizations of the walk ***X***, which satisfies

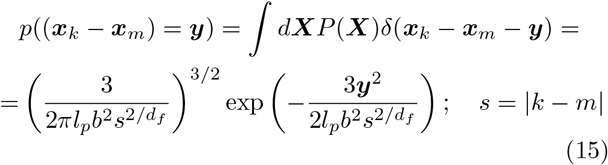

for all *k, m* and some fixed constant *d*_*f*_ called fractal dimension of the walk.

It the recent works of one of us [2, 4, 37] it has been shown that in the limit of large *N* for the subdiffusive (*d*_*f*_ > 2) fractal Brownian motion (fBm) one can write down *P* (***X***) as a Gibbs measure with a pairwise quadratic Hamiltonian:

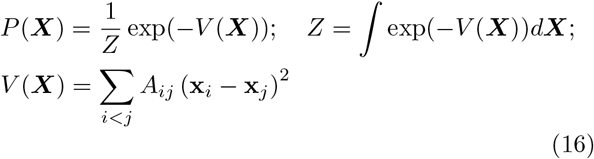

with a proper choice of interactions coefficients *A*_*ij*_ (here *Z* is the partition function, and we use lowercase and uppercase bold letters to denote vectors in three-dimensional and 3 × *N* -dimensional space, respectively). In particular, if the coefficients *A*_*km*_ depend only on the chemical distance between monomers |*k*−*m*| = *s*, so that *A*_*km*_ = *A*(*s*), and if for *s »* 1 *A*(*s*) decays algebraically:

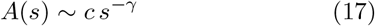

with some *c* > 0, then depending on *γ* there are three possible asymptotic regimes of the chain statistics:

1. If *γ* ≤ 2 all monomers (points of the trajectory) asymptotically merge, and

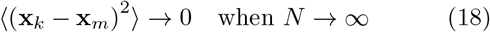

regardless *s* = |*k* − *m*|;
2. If *γ* > 3 the interaction is irrelevant and the large scale properties of the trajectory are indistinguishable from the standard Brownian motion with *d*_*f*_ = 2.
3. Finally, and most interestingly, if 2 < *γ* < 3 the relation (Eq.15) holds for 1 « *s* « *N* with some renormalized 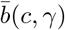 and a non-trivial fractal dimension

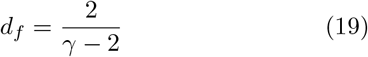

The value of *γ* = 3 is critical, giving rise to the logarithmic corrections to (Eq.15). Note that the conventional beads-on-a-string model corresponds to the choice

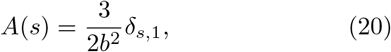

Now we can generalize Eq. (12) to an arbitrary fractal dimension *d*_*f*_ ≥ 2:

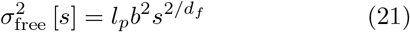

Using the effective Hamiltonian (16) we will generalize Eq. (14) for a fractal bridge in the next section.

#### 3. fBm-bridge

To get the distribution between sites inside a bridge having length *L* (measured in numbers of edges between monomeric units), let’s notice that a fBm-chain becomes a fBm-bridge iff the separation vector between first and last site is **0** (i.e. **x**_0_ = **x**_*L*_). By definition of conditional probability we have for any *s* the following probability distribution function (PDF):

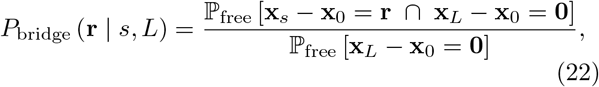

where ℙ_free_ [·] is the probability measure (Eq.16) in a free polymer chain. If chain sections 0…*s* and *s*…*L* were independent, one could represent the numerator as product of two probabilities *P*_free_ (**r** | *s*) and *P*_free_ (**r** | *L* − *s*) and arrive to the Eq. (14) valid for *d*_*f*_ = 2. But a general fBm walk for *d*_*f*_ ≠ 2 has long-range memory, therefore, we need a more delicate technique to express the probability above.

Using the Gibbs distribution Eq. (16) we can write

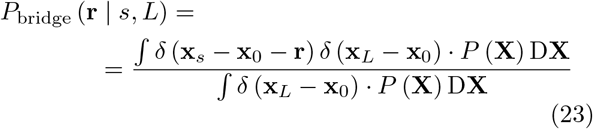

The denominator is calculated straightforward from (Eq.15) and proceeding with the nominator we can Fourier transform the delta functions

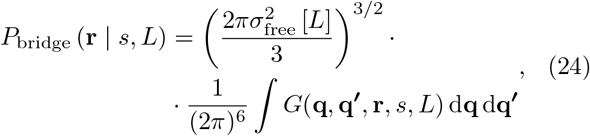

where the Green function has the following form (putting all formulas in (Eq.16) together)

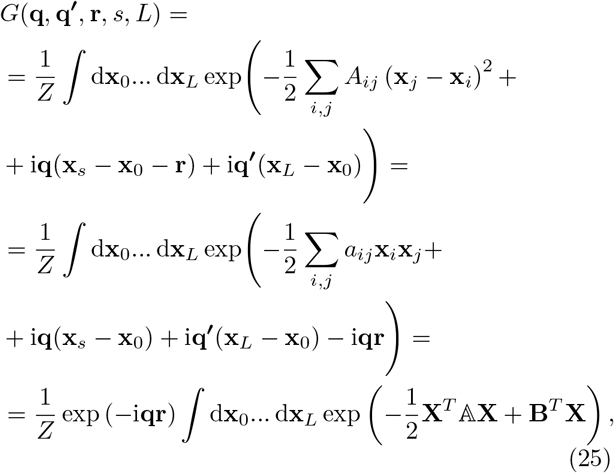

where the new matrix 𝔸 = ‖*a*_*ij*_ ‖ were introduced:

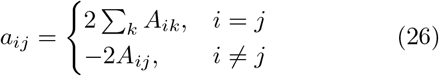

and the new vector **B** ∈ ℝ^3(*L*+1)^ consisted of *L* + 1 parts

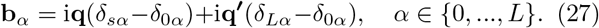

Using new notations we can express the full partition function *Z* (again from (Eq.16))

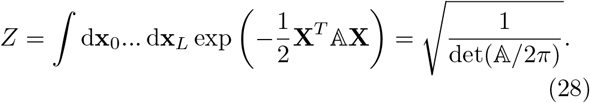

Using the resulting gaussian-like integral in (Eq.25), we get a new form of the Green function:

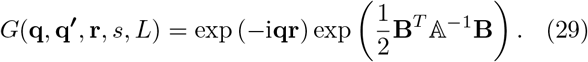

Let us denote *ω*_*p*_ and **A**_*p*_ (*p* = 0, 1, …, *L*) — eigenvalues and eigenvectors of 𝔸, respectively, ordered from the smallest eigenvalue to the largest. Using scalar (inner) products

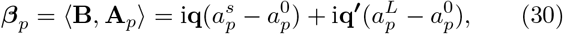

where 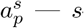-th component of the vector **A**_*p*_, we can express

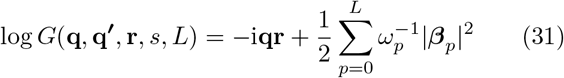

Substituting (Eq.31) into (Eq.24) gives us

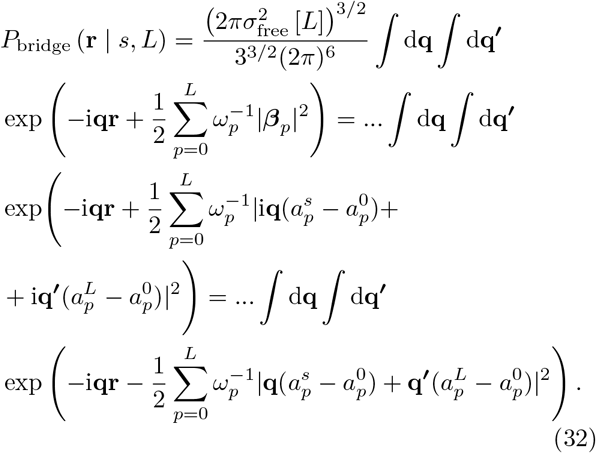

This is a double Gaussian integral, which can be calculated explicitly through parenthesis expansion and grouping components of **q** and **q**^*l*^ into pairs:

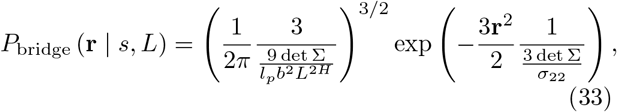

where elements of the matrix Σ are

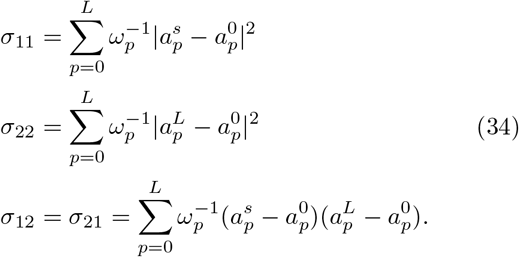

Importantly, the elements of matrix Σ have the meaning of the mean-square distances between corresponding monomers. Indeed, expressing these distances as sums over normal coordinates **u**_*p*_ = ⟨ **X, A**_*p*_⟩ and using the equipartition theorem, which determines the average amplitudes of the normal modes at equilibrium (see [2] for more details), one gets 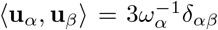 and, therefore,

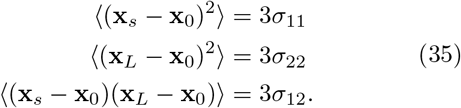

The left hand sides can be easily derived from properties of the fBm:

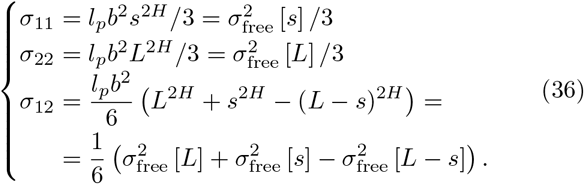

Getting back to Eq. (33) we obtain

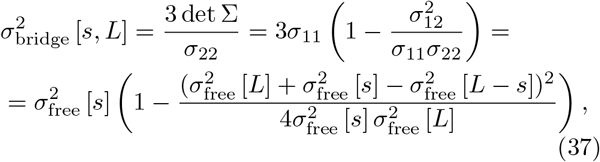

Notice that when *H* = 1/2 (standard Brownian motion)

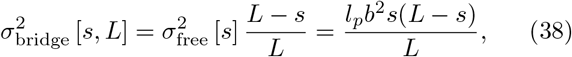

which naturally matches Eq. (14).

#### 4. Conditional contact probabilities for different diagrams

For an arbitrary polymer chain, the pairwise contact probability between two sites separated by linear distance *s* is given by

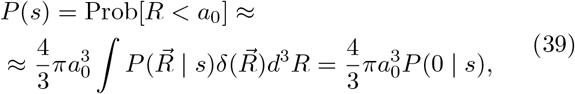

where *a*_0_ is a cutoff contact-radius, and 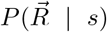 is the probability distribution of the inter-sites separation vector 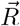. In derivation of Eq. (39) we assumed that 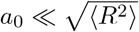.

From Eq. (39) we immediately find that the contact probability between two sites of a fBm chain with fractal dimension *d*_*f*_ and free ends behaves as 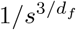, where *s* is the contour separation between the points.

How does this result change in the presence of random loops? For a given realisation of the random loops array, one should consider four scenarios of the relative positions of the two sites as schematically shown in Fig. 7. Here we neglect the diagrams corresponding to the nested and overlapping loops configurations as they rarely occur under the conditions of the interphase. Under the assumption of equilibrium loops (quenched disorder of loops), the resulting contact probability is given by the sum

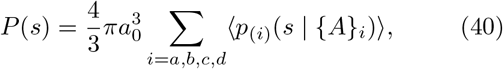

where

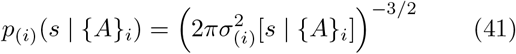

is the contact probability conditional to the particular class of subchain. In what follows we work out exact expressions for 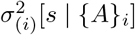.

**FIG. 7.**
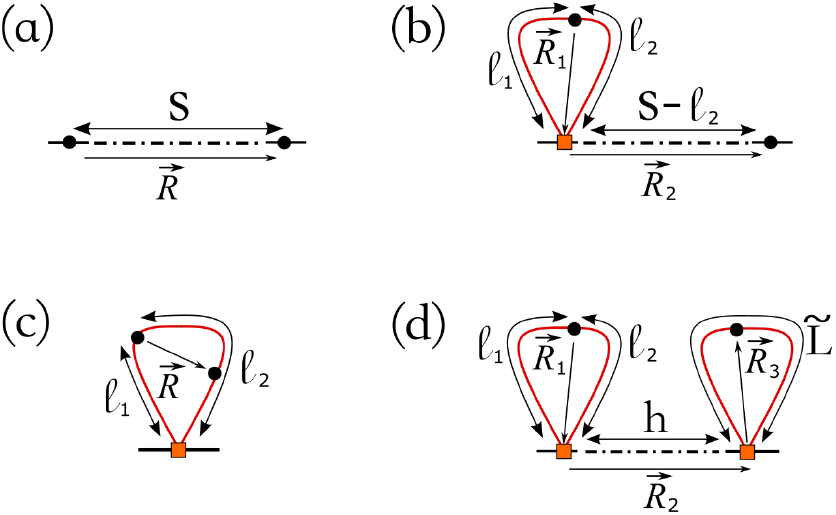
Four diagrams contributing to the contact probability between two sites (depicted as black dots) of polymer chain with a quenched disorder of random loops (depicted in red). The loop extrusion factors are depicted as orange squares.

First, let us consider the diagram (a). In the presence of an arbitrary number of loops between the two points of interest the variance of their physical separation reads

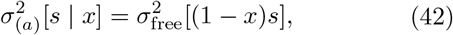

where *x* (0 ≤ *x <* 1) denotes the fraction of the subchain length occupied by the loops. In other words, the intervening loops lead to the effective reduction of the contour distance between the points.

Next, let us consider a subchain of linear size *s* with one end belonging to the gap region and another end belonging to the loop. The loop containing one of two sites of interest can be parameterized by the lengths *l*_1_ and *l*_2_, as shown in Fig 7b. Clearly, the separation vector 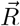 between two sites can be represented as a sum of mutually independent zero mean Gaussian random vectors, 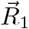 and 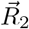, whose statistical properties are described, respectively, by the normal distributions (11) and (13) under an appropriate chose of parameters. Recalling that the sum of independent Gaussian variables has normal statistics with the variance given by the sum of variances of the underlying terms we obtain

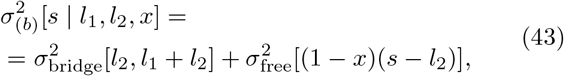

where 0 ≤ *x <* 1, *l*_1_ ≥ 0 and 0 ≤ *l*_2_ ≤ *s*. Now *x* is the fraction of contour length occupied by loops in the segment of size *s*− *l*_2_ enclosed between the loop base and the remaining end of the subchain.

Now let us consider a subchain located inside a loop, see Fig. 7c. Then

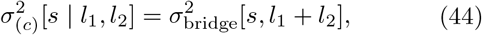

where *l*_1_ ≥ 0 and *l*_2_ ≥ *s*.

Finally, for the last diagram on Fig. 7d

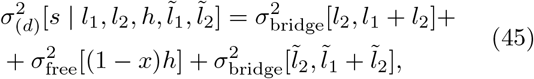

where 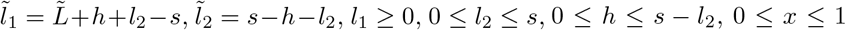 and 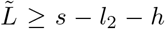. Here *x* is the fraction of contour length occupied by loops in the segment of size 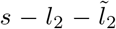 enclosed between two loops depicted in Fig. 7d.

#### 5. Statistical weights of the diagrams

To perform the averaging procedure over disorder of loops assumed in Eq. (40) one needs to know the probability distributions of the random variables {*A*_*i*_}, which parameterize contributions coming form different types of the diagrams (a-d).

Let us treat the sizes of the loops and of the gaps as the independent random variables having exponential probability distributions with *α*_*l*_ = *λ*^−1^ and *α*_*g*_ = *g*^−1^. Then, in order to derive the statistical weights of the diagrams, it is convenient to introduce an auxiliary Markov jump process with two states, “Loop” and “Gap”, in continuous time where time intervals are measured in the units of the polymer contour length. Clearly, the statistics of the random pattern of alternating loops and gaps is analogous to the joint statistics of the time intervals which the Markov process spends in the “Loop” (“L”) and the “Gap” (“G”) states in the course of its stochastic dynamics.

Based on this analogy, we can express the statistical weight of the diagram (*a*) as follows

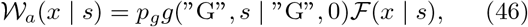

where 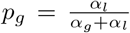 is the probability to find the statistically stationary Markov process in the state “Gap” if you visit it in a random time moment (i.e. the probability that a randomly chosen point of the polymer belongs to a gap), *g*(“G”, *s* | “G”, 0) 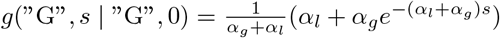 is the probability to find the Markov jump process in the state “Gap” after time *s* under the condition that it starts in the state “Gap” (i.e. the probability that the second point, which is separated by contour distance *s* from the first point, also belongs to a gap region) and ℱ (*x* | *s*) is the probability distribution of the random variable *x* representing the fraction of time that the Markov process spent in the state “Loop” under the condition that it starts in the state “Gap” and find itself in the state “Gap” after the time *s* (i.e., *x* is the fraction of the contour length occupied by loops for a segment whose both ends belong to the gap regions), which is given by (see [39])

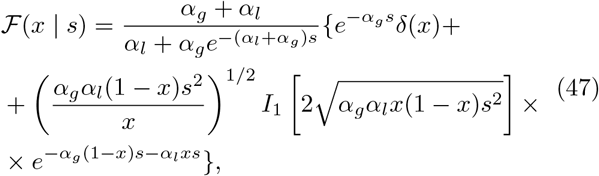

with *I*_1_[*x*] denoting the modified Bessel function of the first kind [56].

Next, the statistical weight of configurations responding to the diagram (*b*) is

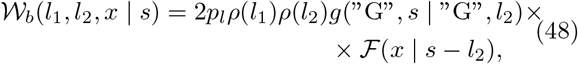

where 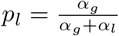 denotes the probability to find the statistically stationary Markov process in the state “Loop” if you visit it in a random time moment, the factors 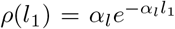 and 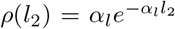 represent, respectively, the probability densities of the random time *l*_1_ elapsed since the last entrance to the state “Loop” and of the random time *l*_1_ remaining before the next jump to the state “Gap”, and ℱ (*x* | *s* −*l*_2_) is the probability distribution of the fraction of time *x* that the Markov process spent in the state “Gap” under the condition that it starts in the state “Gap” and find itself also in the state “Gap” after the time *s* − *l*_2_. Note also that factor 2 in Eq. (48) arises due to the left-right symmetry in the choice of point which is assumed to reside on the loop.

Similarly, the statistical weight of configurations responding to the diagram (*c*) is

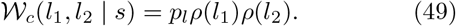

Finally, the statistical weight of subchains described by the diagram (*d*) is given by

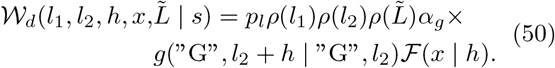

Substituting (42)-(45) and (46)-(50) in Eq. (40) we come to the final expressions of contact probability in the fractal polymer model (see Supplementary Materials).

#### 6. Entanglement length *N*_*e*_

Now let us add to our model of a fractal chain with loops another ingredient, the entanglement length *N*_*e*_.

This parameter allows us to describe the crossover in *r*^2^ between two regimes of *s*:

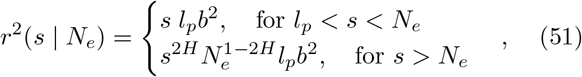

where *H* = 1*/d*_*f*_ and *d*_*f*_ is the fractal dimension. For *d*_*f*_ = 3 this behaviour accounts for the real crumpled polymer chains, such as a ring from the melt of unknotted non-catenated rings [16, 36]. Thus, the physical meaning of *N*_*e*_ is the scale of the minimal crumple, below which the crumpled polymer does not feel topological constraints in the system and has ideal statistics.

Expression (Eq.51) is non-smooth, making the analysis of log-derivatives of the respecting curves problematic. Thus, it is worth to smooth (Eq.51) by averaging over a certain distribution of *N*_*e*_. If we take *N*_*e*_ as an exponentially distributed random variable *x* with mean *N*_*e*_, we can average *r*^2^ above and use it as a new function 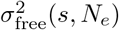 :

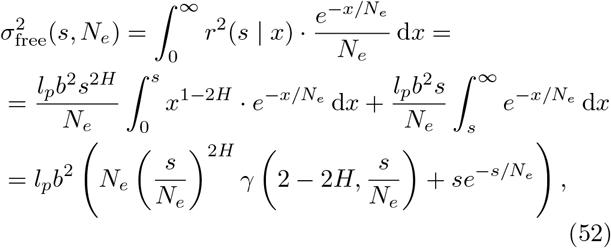

where *γ*(*x, y*) is the lower incomplete gamma function.

Since *l*_*p*_ « *N*_*e*_ one can substitute the lower bound by zero in (Eq.52). The resulting averaged 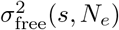 is substituted instead of (21) in Eq. (37) and Eq. (42)-(45). Noticeably, since all bridges and the backbone are independent, we can use different entanglement lengths, one for bridges and one for the backbone.

## APPENDIX B

### DERIVATION OF THE EFFECTIVE VOLUME DENSITY OF A CHAIN FOLDED INTO LOOPS

The increase of the entanglement length for a chain with loops is associated with the decrease of the effective volume density of the polymer, according to the formula (Eq.6). The effective Kuhn volume density 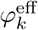 relevant for the entanglements, of a crumpled polymer with loops can be expressed as follows

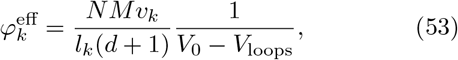

where 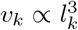 and *l*_*k*_ are the volume and the size of the Kuhn segment, respectively; *M* is the number of chains sharing the volume *V*_0_ of a single chain; *d* = *λ*/*g* is the density parameter (see [38]); 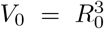 is the volume occupied by one chain, i.e. the cubed size of its backbone; *V*_loops_ is the total volume of the loops. The spatial size of the backbone is controlled by its contour length, *N*_0_ = *N/*(*d* + 1), the entanglement length *N*_*e*_ and the Kuhn length *l*_*k*_ as follows

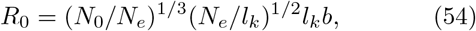

therefore,

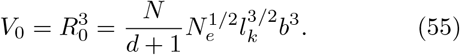

Since each loop is unentangled, their spatial size can be approximated by the size of a Gaussian bridge, 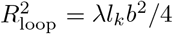, and their overlap with each other can be neglected. The mean number of the loops occupying the volume *V*_0_ reads 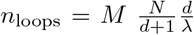. Then one can estimate the total volume of the loops as follows

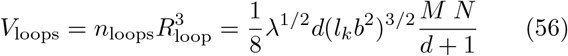

The total number of chains *M* populating the volume *V*_0_ is determined by the total Kuhn volume density of the system 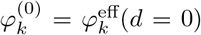 and the entanglement length *N*_*e*_ as

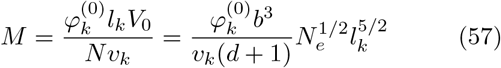

Collecting (Eq.53), (Eq.55), (Eq.56) and (Eq.57) together, one arrives at the following expression for the effective volume density of the backbone

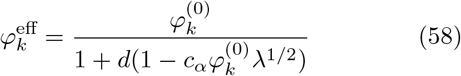

where the constant *c*_*α*_ depends on the parameters of the fiber, 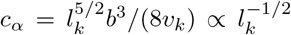. In particular, *c*_*α*_ depends on the numerical coefficients in the expression for the excluded volume of the Kuhn segment, 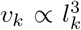, which are determined by the microscopic structure of the fiber. Fitting the resulting 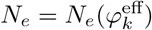 as predicted by (Eq.58) to the one computed in simulations, we find the phenomenological expression for 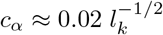.

The structure of (Eq.58) is quite transparent. Without loops (*d* = 0), the backbone resembles the original crumpled chain with effective volume density 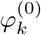. If the loops are sufficiently short so that their volume can be neglected, then the main contribution to the decreased volume density comes from the backbone shortening, which yields the factor *d* + 1 in the denominator. In general case, the loops occupy a certain volume partially compensating the effect of the backbone shortening.

To understand better the role of the compensation effect, it is instructive to introduce the entanglement length 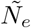 of a *single* (*M* = 1) crumpled chain without loops of the same length *N* and same Kuhn volume density 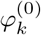.

Clearly from (Eq.57), 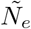 is controlled by the volume density of the chain, 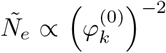.Then the combination 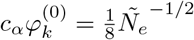 and one can rewrite (Eq.58) as follows

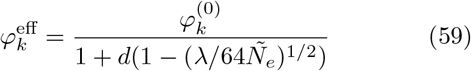

From (Eq.59) it is evident that the compensation effect by the loops is not full, as long as the loops placed on the chain are shorter than its entanglement length (non-topological loops), 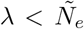. This compensation can be accounted as an effective decrease of the density parameter to *d*_eff_ < *d*, such that

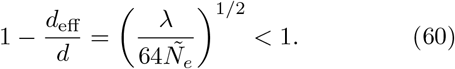

Then, the increase of entanglement length of a chain folded into loops can be described as a result of backbone shortening of a chain with effective density parameter *d*_*eff*_ :

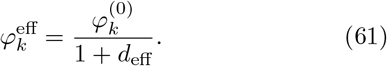

Indeed, for sufficiently short loop sizes 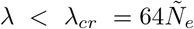, the loops induce the decrease of the effective entanglement density and the associated increase of the entanglement length, *N*_*e*_, of the backbone. At *λ*_*cr*_ the loops are not Gaussian anymore, violating the condition of unentangled loops and, thus, ceasing the dilution of entanglements in the system.

## APPENDIX C

### POLYMER SIMULATIONS

Polymer simulations are done using polychrom (available at https://github.com/open2c/polychrom), a wrapper around the open source GPU-assisted molecular dynamics package OpenMM [57]. In simulations of crumpled polymers with loops the polymer is represented as a closed uknotted chain (ring) of *N* ≈ 90000 monomers with one monomer corresponding to 1 kb, so that the total simulated genomic length is 90Mb. The chains are equipped with harmonic bonds *U*_*bond*_, quadratic angular potential *U*_*angle*_ and excluded volume *U*_*ev*_ interac-tions. Simulations are conducted in the box with periodic boundary conditions (PBC) at volume density *ρΣ*^3^ ≈ 0.3.

The excluded volume potential *U*_*ev*_ is introduced via the auxiliary Weeks-Chandler-Anderson (WCA) potential *U* (*r*_*i*_, *r*_*j*_) [58, 59], which is a lifted Lennard-Jones repulsive branch

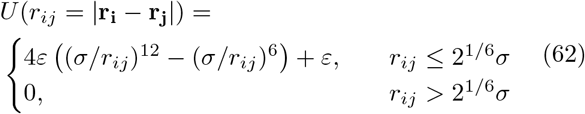

where *Σ* = 0.02*μm* is the characteristic scale of the excluded volume repulsion and *ε* = 1. In order to avoid strong repulsive forces close to the singularity of (Eq.62) in simulations, the WCA potential is further smoothly truncated

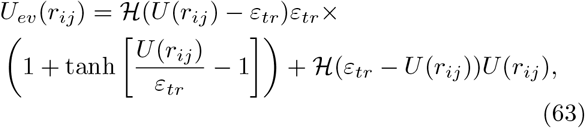

at the prescribed truncation value *ε*_*tr*_ = 10, corresponding to a strong mutual volume exclusion of the beads. In (Eq.63) ℋ is the step function. The potential (Eq.63) acts between every pair of beads, except for neighboring ones.

The energy of harmonic bonds reads as follows

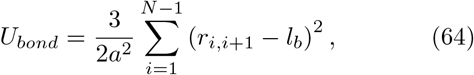

where *a* is the standard deviation of the monomer-to-monomer distance *r*_*i,i*+1_ = |**r**_**i**+**1**_ − **r**_**i**_| from the equilibrium bond length *l*_*b*_. We preset *a* ≈ 0.06*Σ*. In order to repress occasional crossing of two closely located polymer bonds one through another, we choose the value of the equilibrium bond length *l*_*b*_ = 0.8*Σ*. As we found, the value of *l*_*b*_ somewhat smaller than the excluded volume scale facilitates conservation of topology in simulations of crumpled chains.

The angular energy is a harmonic quadratic potential for the consecutive bond angles *θ*_*i,i*+1,*i*+2_ between the corresponding monomers:

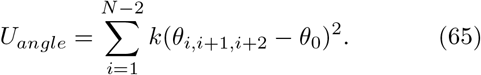

We chose *k* = 2 and *θ*_0_ = *π*. The resulting persistence length of such a model was found to be *l*_*p*_ = 1.75 monomers and the Kuhn segment *l*_*k*_ = 3.5.

In order to test if compartments can interfere with loop extrusion and change the resulting *P* (*s*), we have also introduced block-copolymer interactions in some of our simulations. Annotation into A and B compartments for simulations has been take from Hi-C experiments (the fist eigenvector of the centralised observed over expected HiC map; chromosome 14 in 250kb resolution [60]). The resulting median size of the compartmental domains is ≈ 2.3Mb. The attractive potential between the beads of the same type is the same as described in [61].

First the ring chain without loops is equilibrated in the PBC box starting from a knot-free configuration. After the equilibration the averaged contact probability resembles a typical law for the unknotted non-concatenated rings with 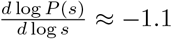 (Fig. S7A).

At the second step we gradually extrude (one-side) the loops on the chain and then fix the loops positions. Extrusion is chosen instead of the instantaneous fixation of the loops in their final positions in order to make sure that the loops are not catenated with each other. We check explicitly that the loops are indeed non-catenated at the end of the extrusion by computing the Gaussian linking number for each pair of the loops. Loops constraints are introduced as additional harmonic bonds between non-neighboring beads of the same energy as the polymer bonds (Eq.64). The loop lengths *l*_1_, *l*_2_, …, *l*_*k*_, *k* = [*N/*(*λ* + *g*)] are drawn randomly from the exponential distribution with the mean *λ* = 100kb. The starting positions of the loop extruders *x*_1_, *x*_2_, …, *x*_*k*_ are drawn randomly, such that the gap lengths *x*_*i*+1_ − *x*_*i*_ − *l*_*i*_ for *i* = 1, 2, …, *k* − 1 are distributed exponentially with the mean *g*. We extrude the loops up to the pre-calculated lengths and then equilibrate the chain with fixed extruders at their final positions for another diffusive relaxation time.

To ensure the chain with loops is sufficiently equilibrated we compute the displacement of a single monomer in the frame of the center-of-mass of the chain (*g*_2_ function in notations [62]) in the course of simulation. As we show in Fig. S7C, at some point the displacements saturate at the gyration size of the whole chain. At this point the monomer starts displacing together with the center-of-mass of the chain (diffusive relaxation time), which is a dynamic evidence of sufficient equilibration of the chain. To provide an estimate of the computation time, such simulation takes ≈ 10 days on a single GPU core (NVidia RTX 2080 Ti).

Quantitative analyses of the chains from simulations (contact probability, *P* (*s*), and end-to-end distances of the backbone chain, 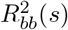 are conducted using the module for the analyses of polymer conformations in polychrom (polymer analyses). The contact probability was computed for two cutoff radii, as shown in Fig. 4D by a strip, cutoff=3*Σ* and 5*Σ*.

To calculate the entanglement length of the backbones in our simulations we have generalized the expression for the end-to-end squared distance *R*^2^(*s*) in the worm-like chain model to account for the crossover at *N*_*e*_. Specifi-cally, we consider the following model

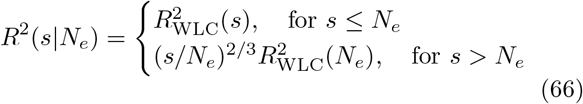

which is further smoothed using the exponential distribution of *N*_*e*_:

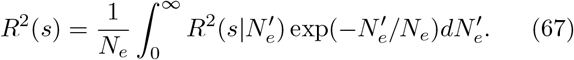

The functional form of 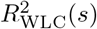 accounts for the tran-sition from the rod-like behavior at short scales to the random-walk behavior at larger scales, describing a persistent polymer chain [48]

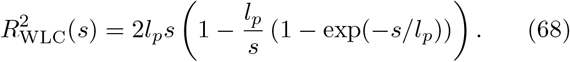

The integration (Eq.67) is performed numerically for various pairs of parameters *l*_*p*_ and *N*_*e*_ with the steps Δ*l*_*p*_ = 0.25 and Δ*N*_*e*_ = 10. The obtained curves are fit to the end-to-end distances computed on the simulated backbone trajectories in the interval [0, 1000] monomers (1Mb), see Figs. S4-S6. The error of the fit *ε*_0_ is computed as the point-wise *L*_2_ distance between theoretical and numerical curves (mean-squared error). The optimal set of (*l*_*p*_, *N*_*e*_) values is determined as the pair yielding the minimum value of the error *ε*_0_. An error of optimal *N*_*e*_ is estimated as the interval around the optimal value yielding the error-of-fit not exceeding 150% of *ε*_0_.

## APPENDIX D

### ESTIMATION OF THE CRITICAL EQUILIBRIUM LOOP EXTRUSION SCALE

The fixed loops model is a reasonable approximation of the active loop extrusion process at scales, for which the extrusion rate *r*_*coh*_ ≈ 1 kb/s is smaller than the rate of thermal relaxation. In contrast with the rate of active extrusion, passive relaxation rate of a polymer non-linearly depends on the contour (genomic) length scale, *s*; e.g., in the framework of the Rouse model, it can be estimated as 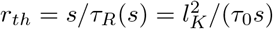, where *τ*_0_ is the microscopic Rouse time (the diffusion time of one monomer at the scale of order of the Kuhn length, [12, 48, 63]). Thus, at short length scales *s < s*^∗^ thermal relaxation is faster than extrusion and the chain is effectively equilibrium, while at larger scales *s > s*^∗^ the non-equilibrium effects should be important. The crossover scale between the two regimes is

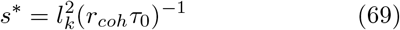

The reported microscopic parameters for chromatin in the literature vary greatly and are quite ambiguous [64, 65]. For the coarse-grained modeling one typ-ically takes *l*_*k*_ = 3 *÷* 4 kb, [66, 67]. As a reference value of the Rouse diffusion coefficient one can take *D* = 10^−2^*μm*^2^*s*^−1/2^, [50, 66]. This yields 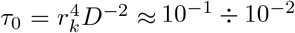 s for the Rouse microscopic time, where we have used the conversion ratio *c* = 60 bp/nm and the Kuhn segment size *r*_*k*_ ≈ 50 nm. Note, however, that a small ≈ 10 nm uncertainty (20%) in the Kuhn segment size translates into an order of magnitude uncertainty for the microscopic Rouse time.

Plugging these values to (Eq.69) one arrives at the following rough estimate of the crossover genomic scale

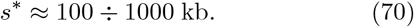

