## Supplemental Information for "Crumpled polymer with loops recapitulates key features of chromosome organization"

### I. INTEGRAL EXPRESSIONS FOR THE CONTRIBUTIONS TO THE CONTACT PROBABILITY

Substituting variances of the squared spatial distance between the points of interest $\sigma^2_{(i)}[s \mid ..]$ and statistical weights $\mathcal{W}_i$ for different diagrams $i = a, b, c, d$ (see Appendix A in the main text), one can arrive at the expressions for the averaged contact probabilities conditional to particular classes of the subchain. Each of the following provide the contributions to Eq. 40 in the Appendix A, being functions of the separation distance $s$ only, while the averaging is performed over the parameters indicated after the "|" sign with the corresponding weight.

$$\langle p_{(a)}(s \mid x)\rangle = \int_0^1 \mathrm{d}x\ \mathcal{W}_a(x \mid s) \left(2\pi\sigma^2_{(a)}[s \mid x]\right)^{-3/2} = \tag{1}$$

$$= \frac{\alpha_l}{\alpha_l + \alpha_g} \cdot \left( e^{-\alpha_g s} \left(2\pi\sigma^2_{(a)}[s \mid 0]\right)^{-3/2} + \int_0^1 \mathrm{d}x\ s \left(\frac{\alpha_l \alpha_g (1-x)}{x}\right)^{1/2} I_1\left[2s\sqrt{\alpha_l \alpha_g x(1-x)}\right] \times \right. \tag{2}$$

$$\left. \times\, e^{-(\alpha_g(1-x)+x\alpha_l)s} \left(2\pi\sigma^2_{(a)}[s \mid x]\right)^{-3/2}\right), \tag{3}$$

$$\langle p_{(b)}(s \mid l_1, l_2, x)\rangle = \int_0^{+\infty} \mathrm{d}l_1 \int_0^s \mathrm{d}l_2 \int_0^1 \mathrm{d}x\ \mathcal{W}_b(l_1, l_2, x \mid s) \left(2\pi\sigma^2_{(b)}[s \mid l_1, l_2, x]\right)^{-3/2} = \tag{4}$$

$$= \frac{2\alpha_g\alpha_l^2}{\alpha_l + \alpha_g} \cdot \left( \int_0^{+\infty} \mathrm{d}l_1 \int_0^s \mathrm{d}l_2\ e^{-\alpha_l(l_1+l_2)-\alpha_g(s-l_2)} \left(2\pi\sigma^2_{(b)}[s \mid l_1, l_2, 0]\right)^{-3/2} + \int_0^{+\infty} \mathrm{d}l_1 \int_0^s \mathrm{d}l_2 \int_0^1 \mathrm{d}x\ (s-l_2) \times \right. \tag{5}$$

$$\left. \times \left(\frac{\alpha_l\alpha_g(1-x)}{x}\right)^{1/2} I_1\left[2(s-l_2)\sqrt{\alpha_l\alpha_g x(1-x)}\right] e^{-\alpha_l(l_1+l_2)-(\alpha_g(1-x)+x\alpha_l)(s-l_2)} \cdot \left(2\pi\sigma^2_{(b)}[s \mid l_1, l_2, x]\right)^{-3/2}\right), \tag{6}$$

$$\langle p_{(c)}(s \mid l_1, l_2)\rangle = \int_0^{+\infty} \mathrm{d}l_1 \int_s^{+\infty} \mathrm{d}l_2\ \mathcal{W}_c(l_1, l_2 \mid s) \left(2\pi\sigma^2_{(c)}[s \mid l_1, l_2]\right)^{-3/2} = \tag{7}$$

$$= \frac{\alpha_g\alpha_l^2}{\alpha_l + \alpha_g} \cdot \int_0^{+\infty} \mathrm{d}l_1 \int_s^{+\infty} \mathrm{d}l_2\ e^{-\alpha_l(l_1+l_2)} \left(2\pi\sigma^2_{(c)}[s \mid l_1, l_2]\right)^{-3/2}, \tag{8}$$

$$\langle p_{(d)}(s \mid l_1, l_2, h, \tilde{L}, x)\rangle = \int_0^{+\infty} \mathrm{d}l_1 \int_0^{s} \mathrm{d}l_2 \int_0^{s-l_2} \mathrm{d}h \int_{s-l_2-h}^{+\infty} \mathrm{d}\tilde{L} \int_0^1 \mathrm{d}x \; \mathcal{W}_d(l_1, l_2, h, x, \tilde{L} \mid s) \left(2\pi\sigma^2_{(d)}[s \mid l_1, l_2, h, \tilde{L}, x]\right)^{-3/2} = \tag{9}$$

$$= \frac{\alpha_g^2 \alpha_l^3}{\alpha_l + \alpha_g} \cdot \left( \int_0^{+\infty} \mathrm{d}l_1 \int_0^{s} \mathrm{d}l_2 \int_0^{s-l_2} \mathrm{d}h \int_{s-l_2-h}^{+\infty} \mathrm{d}\tilde{L} \; e^{-\alpha_l(l_1+l_2) - \alpha_l \tilde{L} - \alpha_g h} \left(2\pi\sigma^2_{(d)}[s \mid l_1, l_2, h, \tilde{L}, 0]\right)^{-3/2} + \right. \tag{10}$$

$$+ \int_0^{+\infty} \mathrm{d}l_1 \int_0^{s} \mathrm{d}l_2 \int_0^{s-l_2} \mathrm{d}h \int_{s-l_2-h}^{+\infty} \mathrm{d}\tilde{L} \int_0^1 \mathrm{d}x h \left(\frac{\alpha_g \alpha_l (1-x)}{x}\right)^{1/2} I_1\left[2h\sqrt{\alpha_g \alpha_l x(1-x)}\right] e^{-\alpha_l(l_1+l_2) - \alpha_l \tilde{L} - \alpha_g(1-x)h - \alpha_l x h} \cdot \tag{11}$$

$$\left. \cdot \left(2\pi\sigma^2_{(d)}[s \mid l_1, l_2, h, \tilde{L}, x]\right)^{-3/2} \right), \tag{12}$$

where

$$\begin{aligned}
\sigma^2_{\mathrm{(a)}}\left[s, x\right] &= \sigma^2_{\mathrm{free}}\left[s(1-x)\right], \\
\sigma^2_{\mathrm{(b)}}\left[s, l_1, l_2, x\right] &= \sigma^2_{\mathrm{bridge}}\left[l_2, l_1 + l_2\right] + \sigma^2_{\mathrm{free}}\left[(s - l_2)(1-x)\right], \\
\sigma^2_{\mathrm{(c)}}\left[s, l_1, l_2\right] &= \sigma^2_{\mathrm{bridge}}\left[s, l_1 + l_2\right], \\
\sigma^2_{\mathrm{(d)}}\left[s, l_1, l_2, h, \tilde{L}, x\right] &= \sigma^2_{\mathrm{bridge}}\left[l_2, l_1 + l_2\right] + \sigma^2_{\mathrm{free}}\left[h(1-x)\right] + \sigma^2_{\mathrm{bridge}}\left[s - h - l_2, \tilde{L}\right]
\end{aligned} \tag{13}$$

and

$$\sigma^2_{\mathrm{free}}\left[s\right] = l_p b^2 s^{2/d_f} \tag{14}$$

for a fractal polymer with arbitrary fractal dimension $d_f \geq 2$ and

$$\sigma^2_{\mathrm{free}}\left[s\right] = l_p b^2 \left( N_e \left(\frac{s}{N_e}\right)^{2H} \gamma\left(2 - 2H, \frac{s}{N_e}\right) + s e^{-s/N_e} \right), \tag{15}$$

for a crumpled chain (non-fractal) with exponentially distributed entanglement length with mean $N_e$. In both cases

$$\sigma^2_{\mathrm{bridge}}\left[s, L\right] = \sigma^2_{\mathrm{free}}\left[s\right] \left( 1 - \frac{\left(\sigma^2_{\mathrm{free}}\left[L\right] + \sigma^2_{\mathrm{free}}\left[s\right] - \sigma^2_{\mathrm{free}}\left[L - s\right]\right)^2}{4\sigma^2_{\mathrm{free}}\left[s\right]\sigma^2_{\mathrm{free}}\left[L\right]} \right) \tag{16}$$

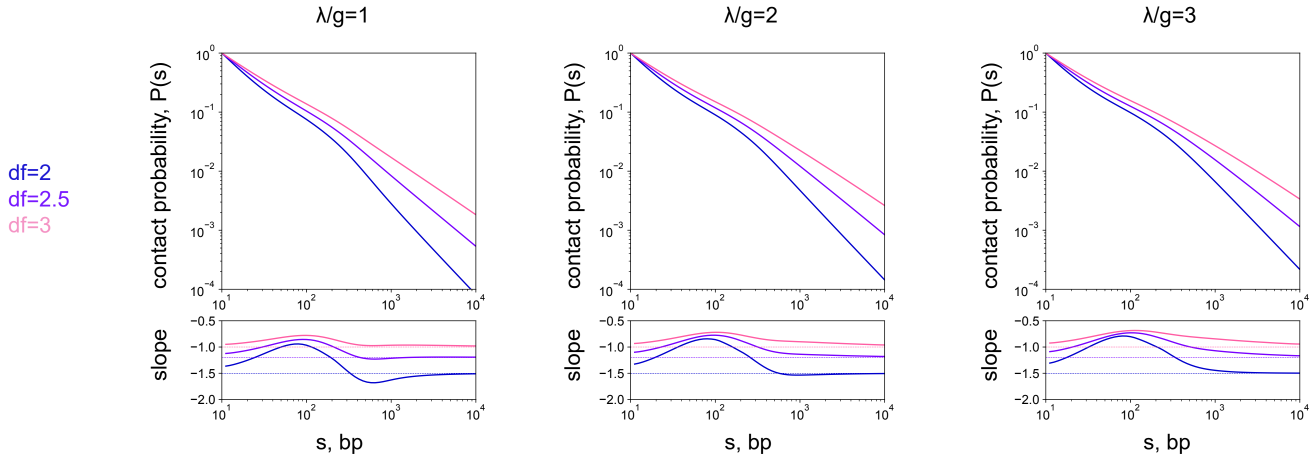


FIG. S1. The contact probability for fractal chains with various fractal dimensions $d_f$ for various values of the parameter $\lambda/g$ (the mean loop size is fixed to $\lambda = 100$kb). Critical value of the density parameter, at which the dip disappears: $(\lambda/g)^* \approx 2$ (for $d_f = 2$), $(\lambda/g)^* \approx 1$ (for $d_f = 3$).

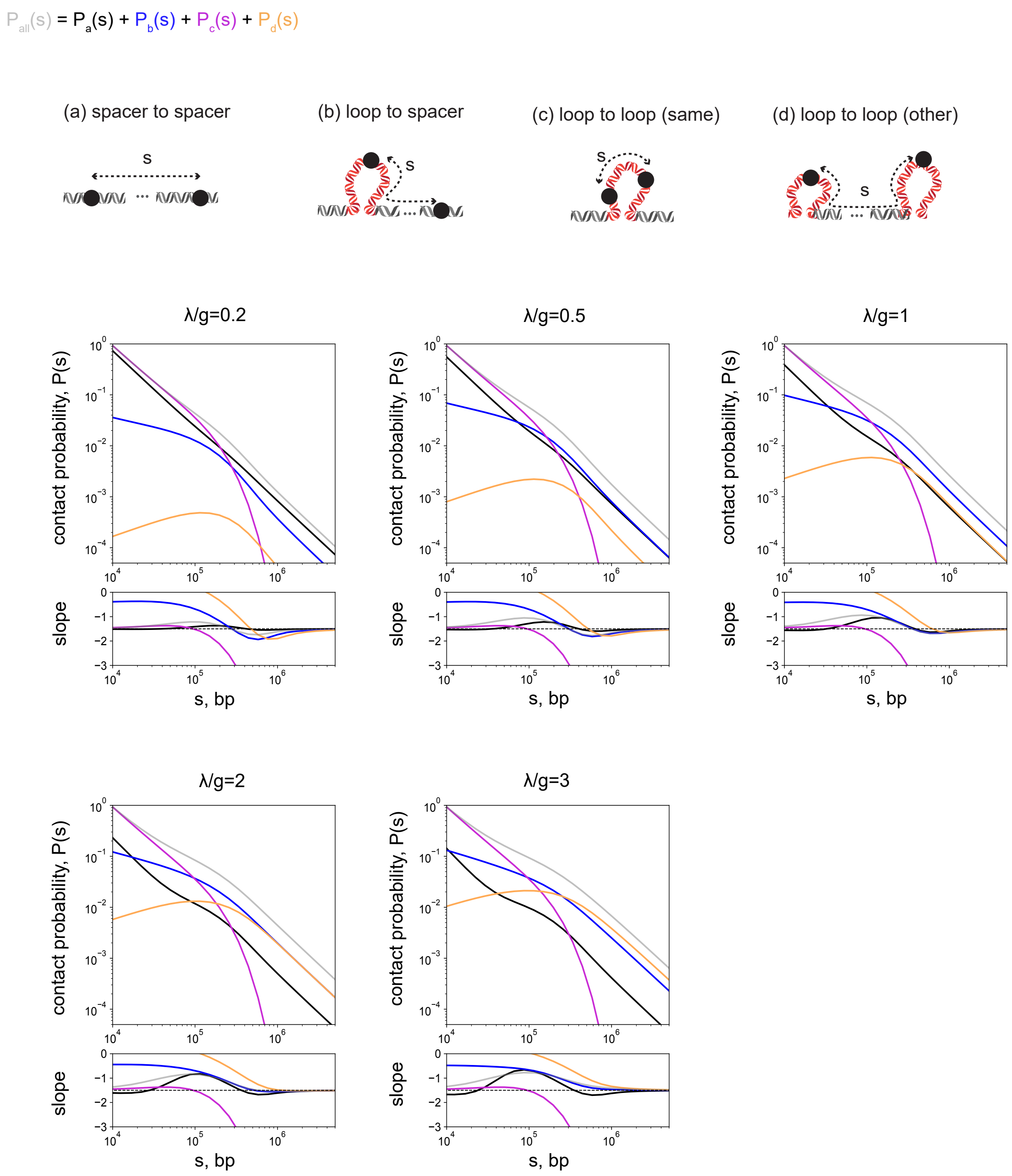


FIG. S2. Contributions of different diagrams to the contact probability at various $\lambda/g$. The mean loop size $\lambda = 100$kb, fractal dimension $d_f = 2$. $P_a$: contacts between spacers; $P_b$: contacts between loop and spacer; $P_c$: contacts within one loop; $P_d$: contacts between different loops. Refer to Figure 2A for schematics.

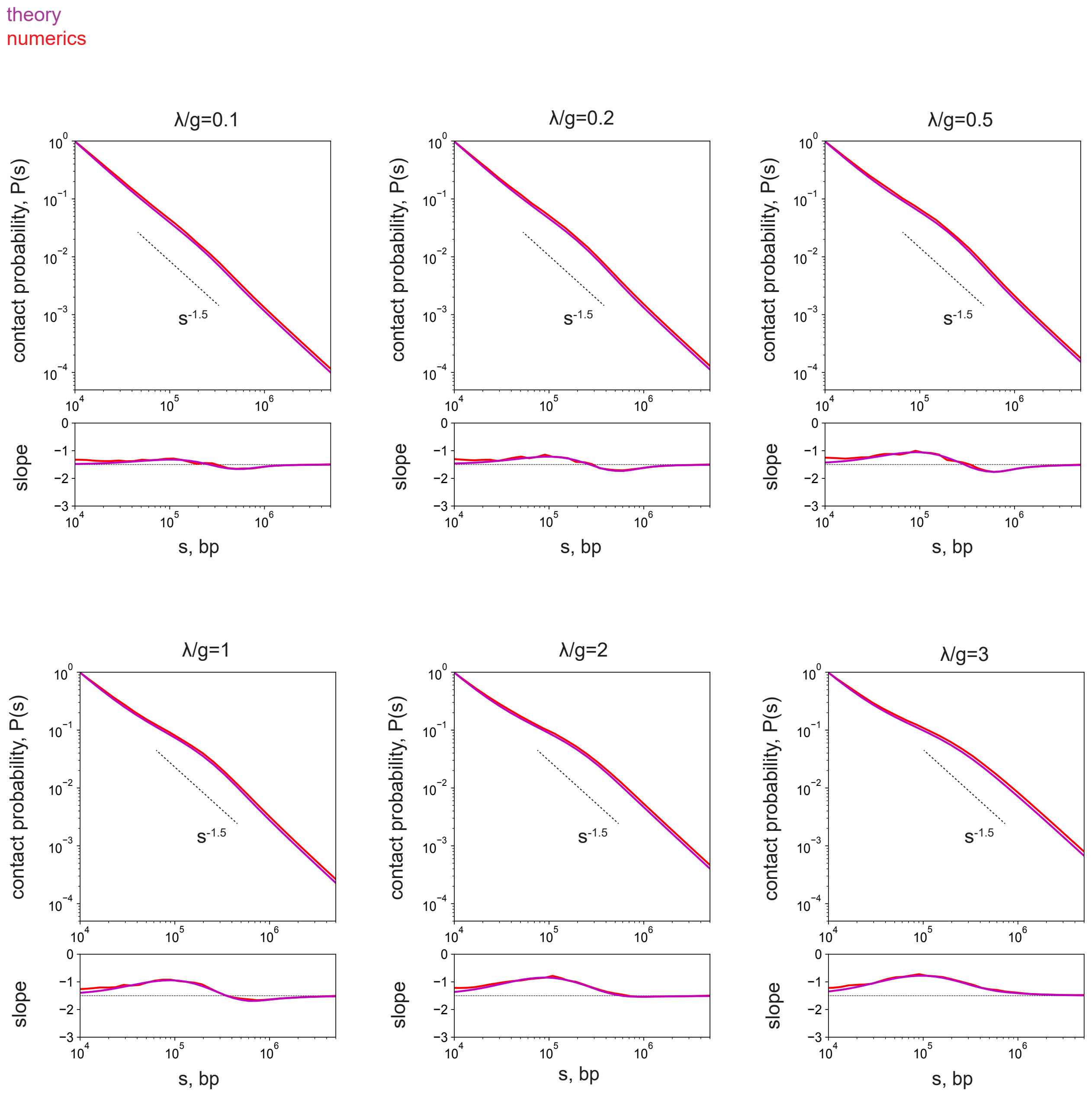


FIG. S3. Full theory vs. numerics for various values of $\lambda/g$ ($\lambda = 100\text{kb}, d_f = 2$).

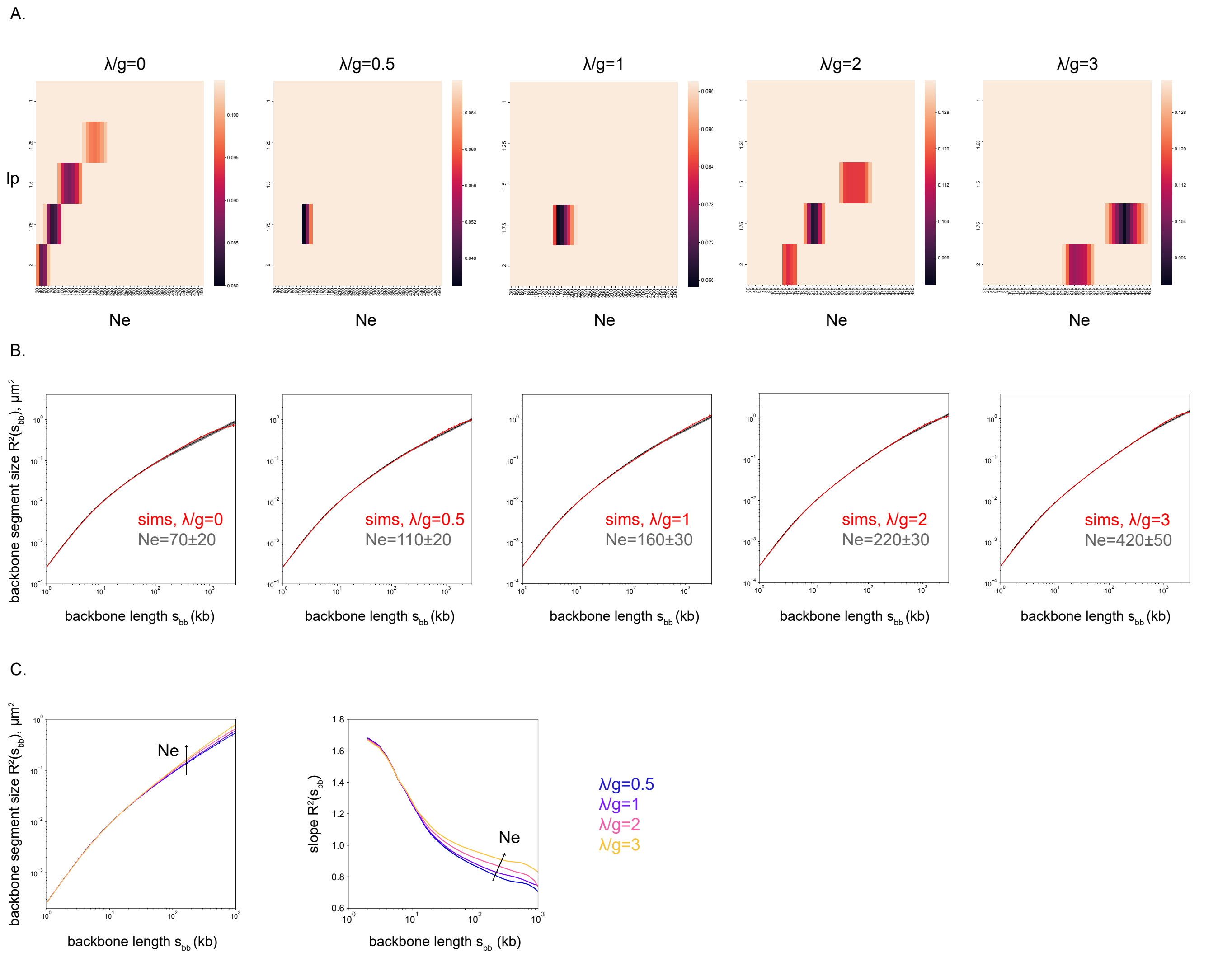


FIG. S4. (A). Heatmaps of the mean squared error (MSE) of the backbone segment size fitting by different $l_p$ and $N_e$ using the extension of the worm-like chain model for crumpled chains (see Appendix C). The range of the colorbar corresponds to the $[\varepsilon_0; 1.5\ \varepsilon_0]$, where $\varepsilon_0$ is the minimal error for each $\lambda/g$. (B). The corresponding end-to-end squared distances $R^2(s_{bb})$ for the backbone segments $s_{bb}$. Simulations curves are shown in red, the best-fit theoretical curves by dashed black and the error is shown by thick gray curves. (C). The squared sizes of the backbone segments for different $\lambda/g$ (left). The respective log-derivatives (right).

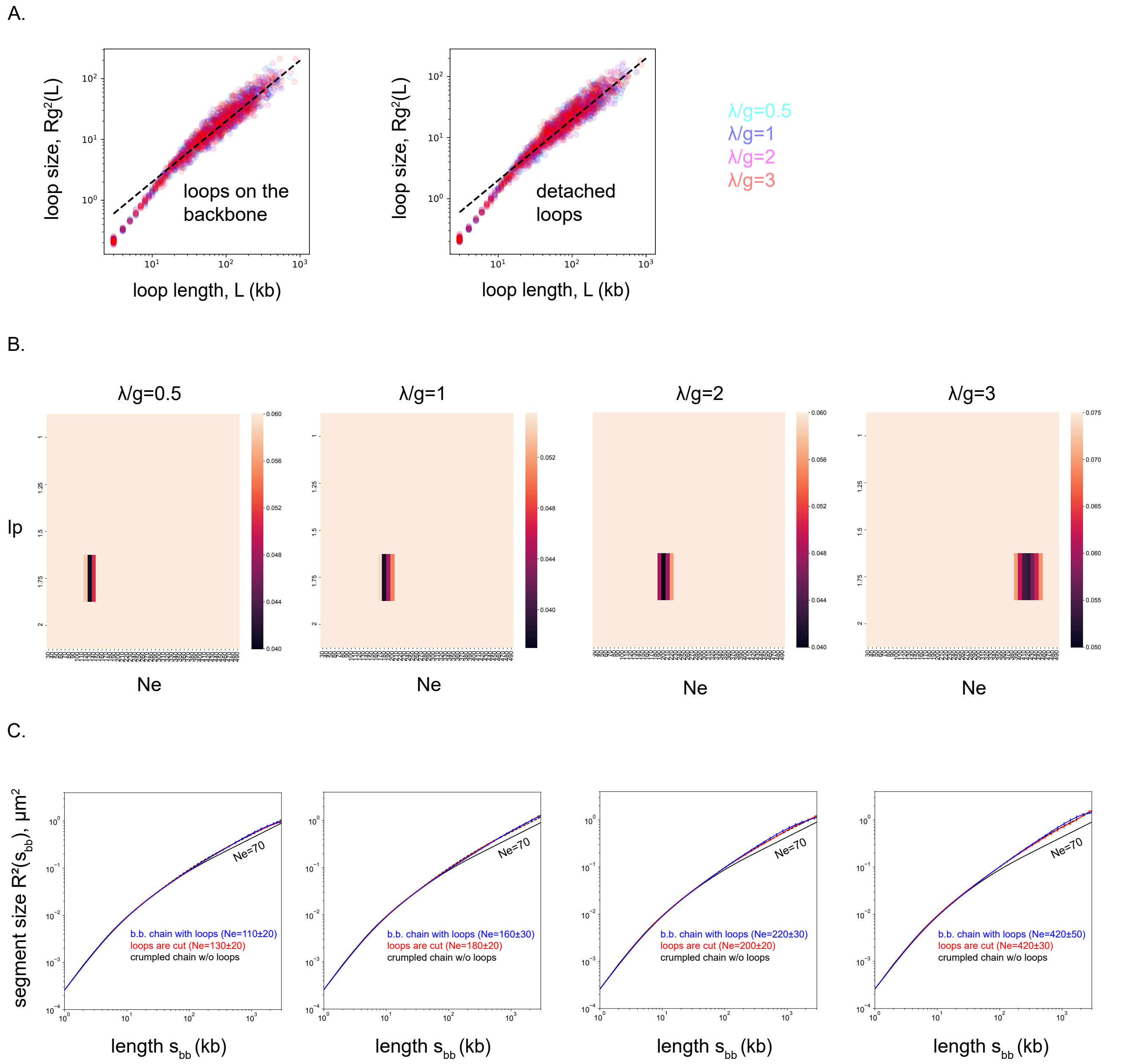


FIG. S5. Simulations of the chain with detached (cut) loops. (A). Squared sizes of the loops for the chain folded into loops (left). The same for a chain with disconnected loops from it (right). (B). Heatmaps of the mean squared error (MSE) of the main chain segment size fitting (same as Fig. S4A). (C). The corresponding end-to-end squared distances $R^2(s_{bb})$ for the main chain segments $s_{bb}$ (in red). The best-fit theoretical curve is shown in dashed black. For comparison the backbone segment sizes from Fig. S4B are shown in blue and the theoretical curve corresponding to $N_e = 70$ (a crumpled chain without loops at the same volume density) is shown in solid black.

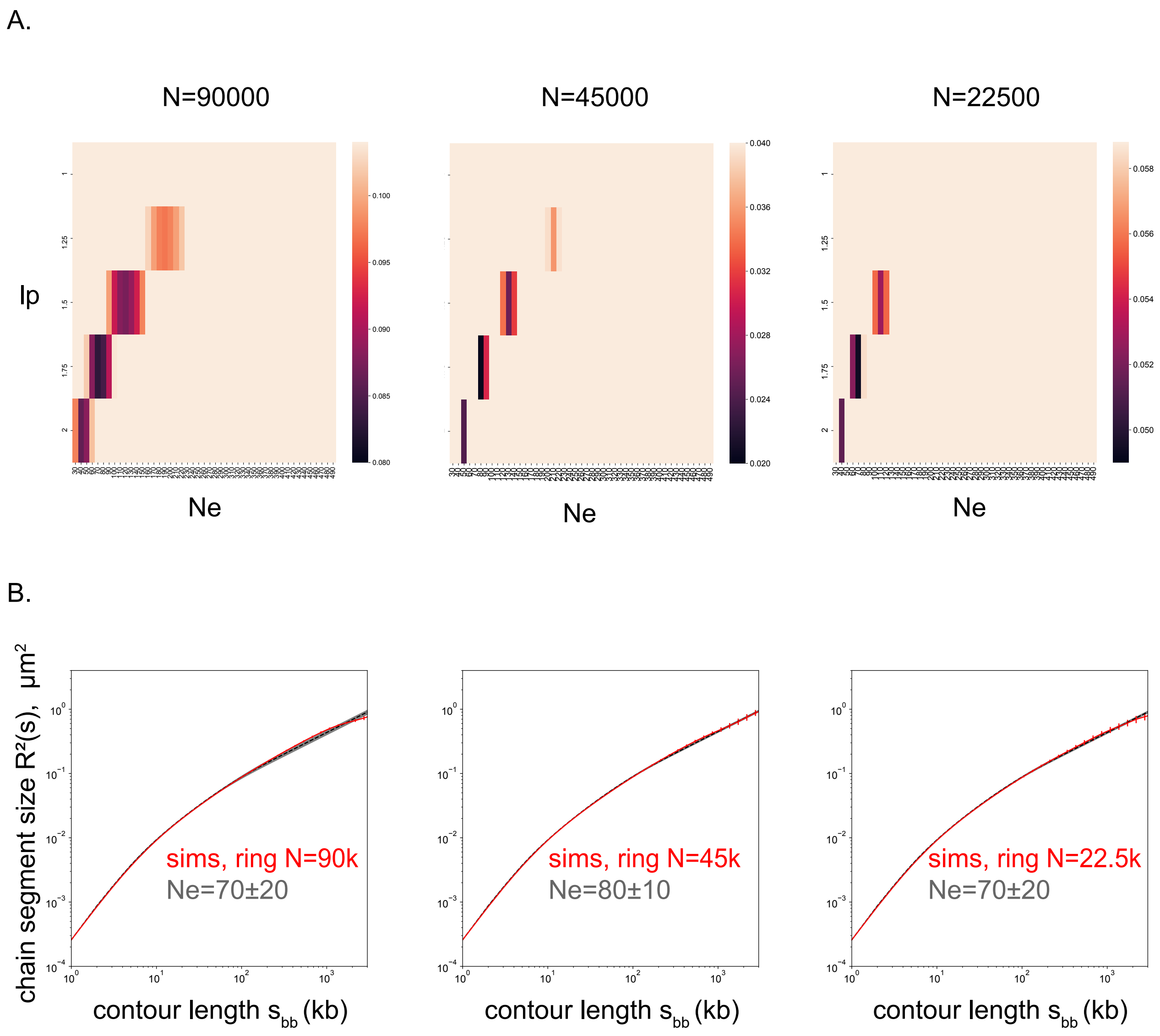


FIG. S6. Simulations of the main chain with removed loops and confined to the same volume density $\varphi \approx 0.3$. (A). Heatmaps of the mean squared error (MSE) of the main chain segment size fitting (same as Fig. S4A and Fig. S5B) for backbones of various length: $N \approx 90000$ ($\lambda/g = 0$), $N \approx 45000$ ($\lambda/g = 1$), $N \approx 22500$ ($\lambda/g = 3$). (B). The corresponding end-to-end squared distances $R^2(s_{bb})$ for the main chain segments $s_{bb}$ (in red). The best-fit theoretical curve is shown in dashed black, the error is shown by thick gray curves.

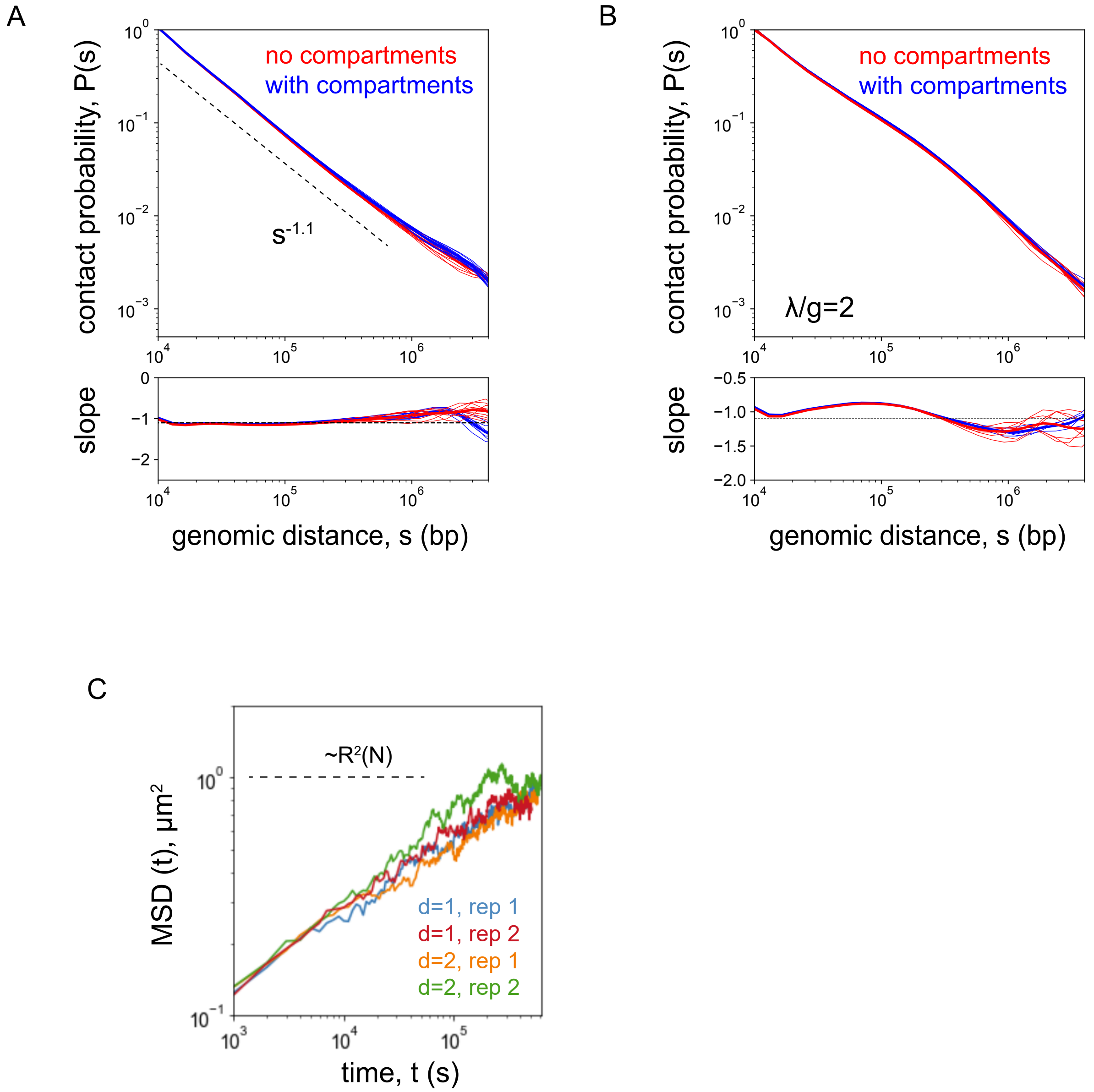


FIG. S7. Simulations. (A). The contact probability function after equilibration of an unknot ring $N = 90$Mb without loops; with (blue) and without (red) compartments. Five consecutive time averages of $P(s)$ for each set is shown. The scaling $s^{-1.1}$, corresponding to crumpled chain with the fractal dimension $d_f = 3$, is not sensitive to the addition of compartmental interactions. The reason is prosaic: the scale of compartmental domains is $\approx$ 2Mb in humans, while the loops are only $\approx 100-200$kb. (B). A similar comparison as in (A) but for a crumpled chain with loops with the density parameter $\lambda/g = 2$. The blue and red set of curves are identical within the statistical error between the averages. (C). The mean-squared displacement of a monomer in the course of equilibration of a chain with loops. Two replicates for two values of $d = \lambda/g$ are shown. The monomer displacement reaches the size of the chain, which is the dynamic evidence of sufficient equilibration of the system. The conversion of timescales from simulations to seconds is done by assuming the value $D \approx 3 * 10^{-2}\mu^2 s^{-1/2}$ of the Rouse diffusion coefficient.

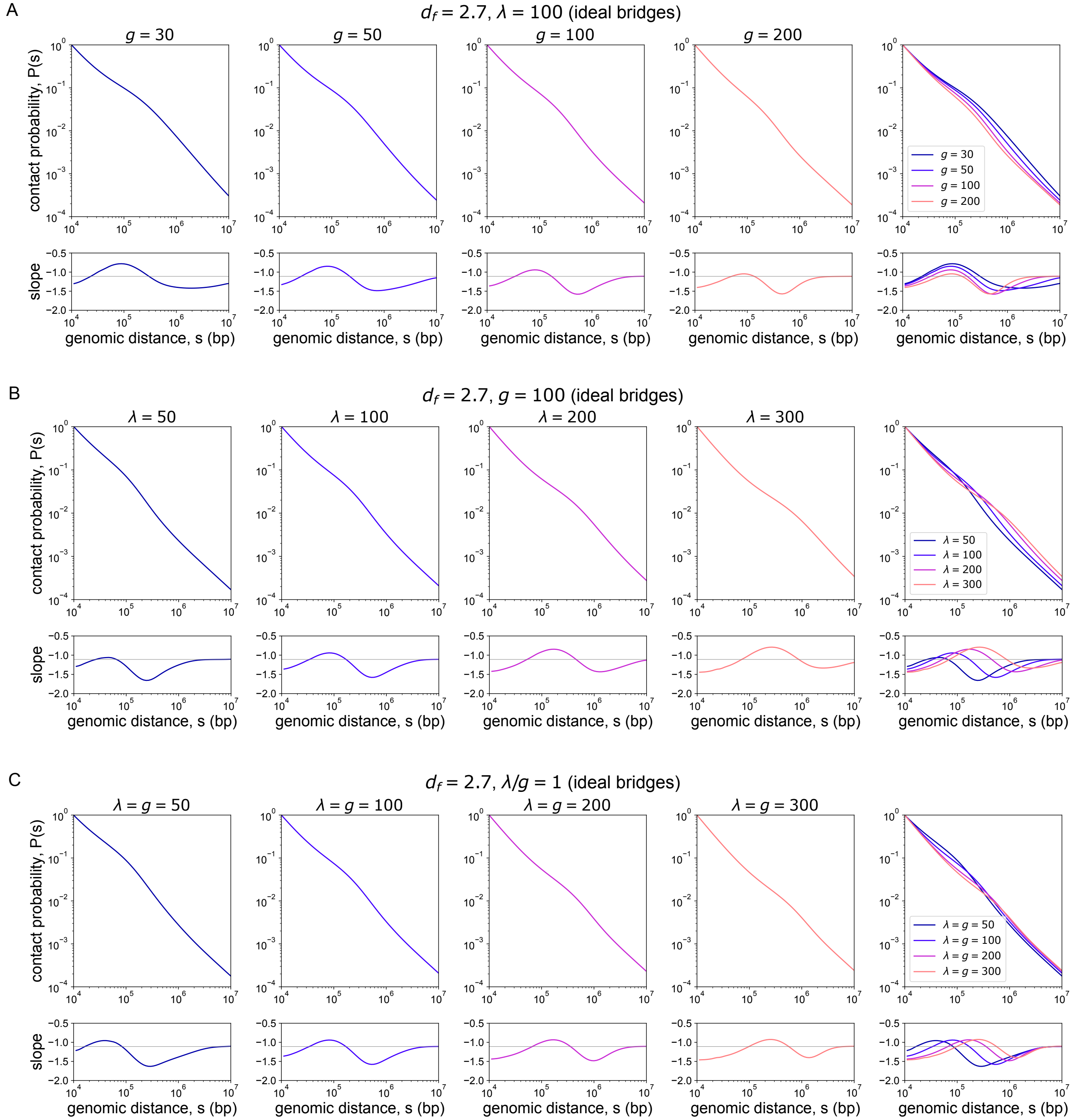


FIG. S8. Change of the contact probability, $P(s)$, in the full theory with ideal loops (Brownian bridges) and entanglement length of the backbone $N_e = N_e(\lambda, d)$. Fractal dimension is $d_f = 2.7$, which corresponds to the slope $\approx -1.1$ of $P(s)$ without loops in the Gaussian model. Volume density $\varphi = 0.2$ and Kuhn length $l_k = 3.5$kb are fixed. (A). Change of spacer size with $\lambda = 100$kb. (B) Change of loop size with $g = 100$kb. (C). Change of loop and spacer sizes accordingly at the constant density parameter $d = \lambda/g = 1$.
